# Heterogeneous loop model to infer 3D chromosome structures from Hi-C

**DOI:** 10.1101/574970

**Authors:** Lei Liu, Min Hyeok Kim, Changbong Hyeon

## Abstract

Adapting a well-established formalism in polymer physics, we develop a minimalist approach to infer threedimensional (3D) folding of chromatin from Hi-C data. The 3D chromosome structures generated from our heterogeneous loop model (HLM) are used to visualize chromosome organizations that can substantiate the measurements from FISH, ChIA-PET, and RNA-Seq signals. We demonstrate the utility of HLM with several case studies. Specifically, the HLM-generated chromosome structures, which reproduce the spatial distribution of topologically associated domains (TADs) from FISH measurement, show the phase segregation between two types of TADs explicitly. We discuss the origin of cell-type dependent gene expression level by modeling the chromatin globules of *α*-globin and SOX2 gene loci for two different cell lines. We also use HLM to discuss how the chromatin folding and gene expression level of Pax6 loci, associated with mouse neural development, is modulated by interactions with two enhancers. Finally, HLM-generated structures of chromosome 19 of mouse embryonic stem cells (mESCs), based on single-cell Hi-C data collected over each cell cycle phase, visualize changes in chromosome conformation along the cell cycle. Given a contact frequency map between chromatic loci supplied from Hi-C, HLM is a computationally efficient and versatile modeling tool to generate chromosome structures, which can complement interpreting other experimental data.

## INTRODUCTION

Recent advances in chromosome conformation capture techniques combined with parallel sequencing^1–5^ and fluorescence imaging microscopies have ushered in a new era of chromosome research over the past decade. Along with post-translational histone modifications, which have been led to conceptualization of epigenomes^6^, the critical findings from fluorescence imaging and Hi-C data, that the spatial organization of chromatin varies with the tissue or cell types^7, 8^, cell cycle^4^, and pathological states^9–11^, have brought a new dimension to our understanding of genome functions.

Among others, maps of genome-wide contact frequencies, quantified by Hi-C data, offer unprecedented opportunities to infer 3D chromosome structures in cell nuclei^12–22^. In a nutshell, Hi-C provides the contact frequencies of genomic loci pairs based on the statistics of PCR-amplified DNA fragments digested from formaldehyde cross-linked cells^1, 2^. One could interpret that Hi-C measures the population-sampled contact probability between pair of genomic loci, say *i* and *j, p_ij_*. A proper mathematical mapping of *p_ij_* to the spatial distance *r_ij_* is of critical importance for interpreting fluorescence imaging data^23, 24^ in comparison with Hi-C data.

The advent of fluorescence *in situ* hybridization (FISH) followed by C-based techniques have engendered much devotion to capture the principle underlying the three-dimensional (3D) folding of chromosomes. This has led to development of a series of polymer-based models over the decades, which include “multiloop subcompartment model,”^25, 26^ “random loop model,” (RLM)^27–29^ “strings and binders switch” model^12, 15, 30^ and its derivative^17, 31, 32^, “loop extrusion model,”^13–15, 33^ “minimal chromatin model,”^34^ and more recently “chromosome copolymer model.”^22^ Among them, while applicability is limited to the associated spatio-temporal scale of the model being considered, some were developed by keeping a specific molecular mechanism in mind or by incorporating “one-dimensional” information of epigenetic modification and/or DNA accessibility along genomic loci as input to heteropolymer model^22, 32, 35^. On the other hand, partly sacrificing the model simplicity, others were developed solely for the purpose of reconstructing more precise 3D chromatin structures from Hi-C^20, 36–38^ and other experiments^39^.

As the cell imaging data over different cell types is rapidly growing, comparative study of chromosome conformations has become imperative. In the abovementioned models, however, a physically sound mapping of *p_ij_* from Hi-C to the spatial distance *r_ij_* (see review^40^) is still lacking, and computational cost are still high. To this end, here we develop a minimalist model that allows us to generate chromatin conformations from Hi-C data in a most efficient way and to study the structural characteristics of chromosome at a length scale of interest corresponding to the resolution of the given data. In order to achieve such a goal in a most simplifying manner, one could learn much from literature of generic polymer problems, such as the collapse transition of an isolated polymer chain or macromolecular networks with increasing number of internal bonds^41–44^, and polymer conformation and dynamics inside confinement^45, 46^.

Pushing the polymer physics idea to its extreme, we propose a minimalist approach, termed the heterogeneous loop model (HLM), that allows us to build 3D structures of chromosomes from Hi-C data. HLM adapts the random loop model (RLM) which was originally developed based on a randomly crosslinked polymer chain^27, 28, 49^. In RLM, which represents chromosome conformation in terms of the sum of harmonic potentials, pairwise contact probabilities are expressed analytically in terms of a few model parameters. Here, without sacrificing the mathematical tractability and simplicity of the RLM, we extend the RLM to HLM by allowing the loop interactions to be non-uniform and heterogeneous, such that the resulting loop interactions can best represent a given Hi-C data.

In this study, we apply HLM to various regions of human and mouse genomes that span 1 – 100 Mb at 5 – 500 kb resolution, and generate the corresponding conformational ensemble of chromsomes. We demonstrate the utilities of HLM by comparing the structural information extracted from HLM-generated chromosome ensemble with those implicated from the measurements from FISH^23, 24, 28^, chromatin interaction analysis by paired-end tag sequencing (ChIA-PET)^50, 51^, and previous modeling studies^28, 32, 37, 52, 53^. Through multiple examples this study will demonstrate that HLM is an excellent approach to infer 3D structures from Hi-C data.

## RESULTS

HLM is effectively a multi-block copolymer model in which monomer-monomner interactions (loops) are harmonically restrained with varying interaction strengths (*k_ij_*) (Methods and SI). Mapping the pairwise contact probabilities *p_ij_* from Hi-C to the model parameters *k_ij_* is the essence of HLM. By incorporating a standard Lennard-Jones non-bonded potential slightly below the *θ*-condition, which takes into account the short-range excluded volume interaction between monomers as well as global thermodynamic driving force that induces microphase separation between different monomer types, HLM allows us to generate a conformational ensemble of chromosome structures that reproduces a contact probability matrix that displays close resemblance to an original input Hi-C data. We used HLM to model various genomic regions (see Table I). HLM-generated chromosome conformations were used to interpret the currently available experimental results.

**TABLE I.** Genomic regions simulated in this work.

| Species | Cell line | Hi-C exp. | Chr. | Start(bp) | End(bp) | Resolution (kb) | $N$ | PC<sup>a</sup> | Time (min)<sup>c</sup> | Fig. |
| --- | --- | --- | --- | --- | --- | --- | --- | --- | --- | --- |
| Human | GM12878 | Ref.<sup>3</sup> | chr5 | 90,000,000 | 100,000,000 | 50 | 200 | 0.96 | 4.8 | 6 |
|  | IMR90 | Ref.<sup>3</sup> | chr21 | 14,000,000 | 48,000,000 | 250 | 137 | 0.97 | 0.8 | 1 |
|  | IMR90 | Ref.<sup>3</sup> | chr11 | 59,000,000 | 94,000,000 | 250 | 140 | 0.98 | 1.7 | S3 |
|  | IMR90 | Ref.<sup>3</sup> | chr1 | 150,000,000 | 180,000,000 | 250 | 120 | 0.98 | 0.8 | S4 |
|  | K562 | Ref.<sup>3</sup> | chr16 | 60,000 | 560,000 | 5 | 100 | 0.94 | 0.2 | 2 |
|  | GM12878 | Ref.<sup>3</sup> | chr16 | 60,000 | 560,000 | 5 | 100 | 0.92 | 0.4 | 2 |
|  | hESC | Ref.<sup>47</sup> | chr3 | 179,000 | 184,000 | 40 | 125 | 0.94 | 1.4 | 3 |
|  | HUVEC | Ref.<sup>3</sup> | chr3 | 179,000 | 184,000 | 40 | 125 | 0.95 | 1.8 | 3 |
| Mouse | mESC | Ref.<sup>48</sup> | chr2 | 105,000,000 | 106,000,000 | 8 | 125 | 0.94 | 1.4 | 4 |
|  | NPC | Ref.<sup>48</sup> | chr2 | 105,000,000 | 106,000,000 | 8 | 125 | 0.96 | 1.2 | 4 |
|  | CN | Ref.<sup>48</sup> | chr2 | 105,000,000 | 106,000,000 | 8 | 125 | 0.97 | 1.3 | 4 |
|  | ncx_NPC | Ref.<sup>48</sup> | chr2 | 105,000,000 | 106,000,000 | 8 | 125 | 0.97 | 0.8 | 4 |
|  | ncx_CN | Ref.<sup>48</sup> | chr2 | 105,000,000 | 106,000,000 | 8 | 125 | 0.97 | 1.1 | 4 |
|  | mESC | Ref.<sup>4</sup> | chr19 | 1 | 61,342,430 | 500 | 117 | 0.92<sup>b</sup> | 1.4 | 5 |

<sup>a</sup>The similarity between contact probabilities ( $p_{ij}$ ) from Hi-C and those from modeling is quantified by the Pearson correlation (PC) (see also discussions in SI). <sup>b</sup>From the post-M to pre-M phase, Pearson correlation of mESCs is 0.77, 0.96, 0.96, 0.96, 0.97 and 0.91, respectively. <sup>c</sup>It takes a few minutes to determine the interaction strength parameters by the constrained optimization, namely, obtaining $\tilde{\mathcal{K}}$ from $\mathcal{P}$ .

### Spatial distribution of TADs inferred from HLM in comparison with FISH measurement

Intra-chromosomal distances between TADs in human IMR90 cells, measured by Wang *et al*. through a multiplexed FISH method^23^, have been used as a benchmark for different models^38^. To show the utility of HLM, we model 34 Mb genomic region on chr21 of IMR90 cells, which contains 33 labeled TADs (Table S1 provides the genomic positions of these TADs).

First, the contact probability matrix 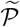 constructed from HLM-generated structures captures the characteristic checkerboard pattern of the heatmap of Hi-C data, 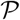; the mean contact probability *P*_HLM_(*s*) of HLM is consistent with *P*_Hi-C_(*s*) calculated from Hi-C over all length scales including the wiggly pattern at large *s* (Figs. 1A and 1B).

**FIG. 1.**
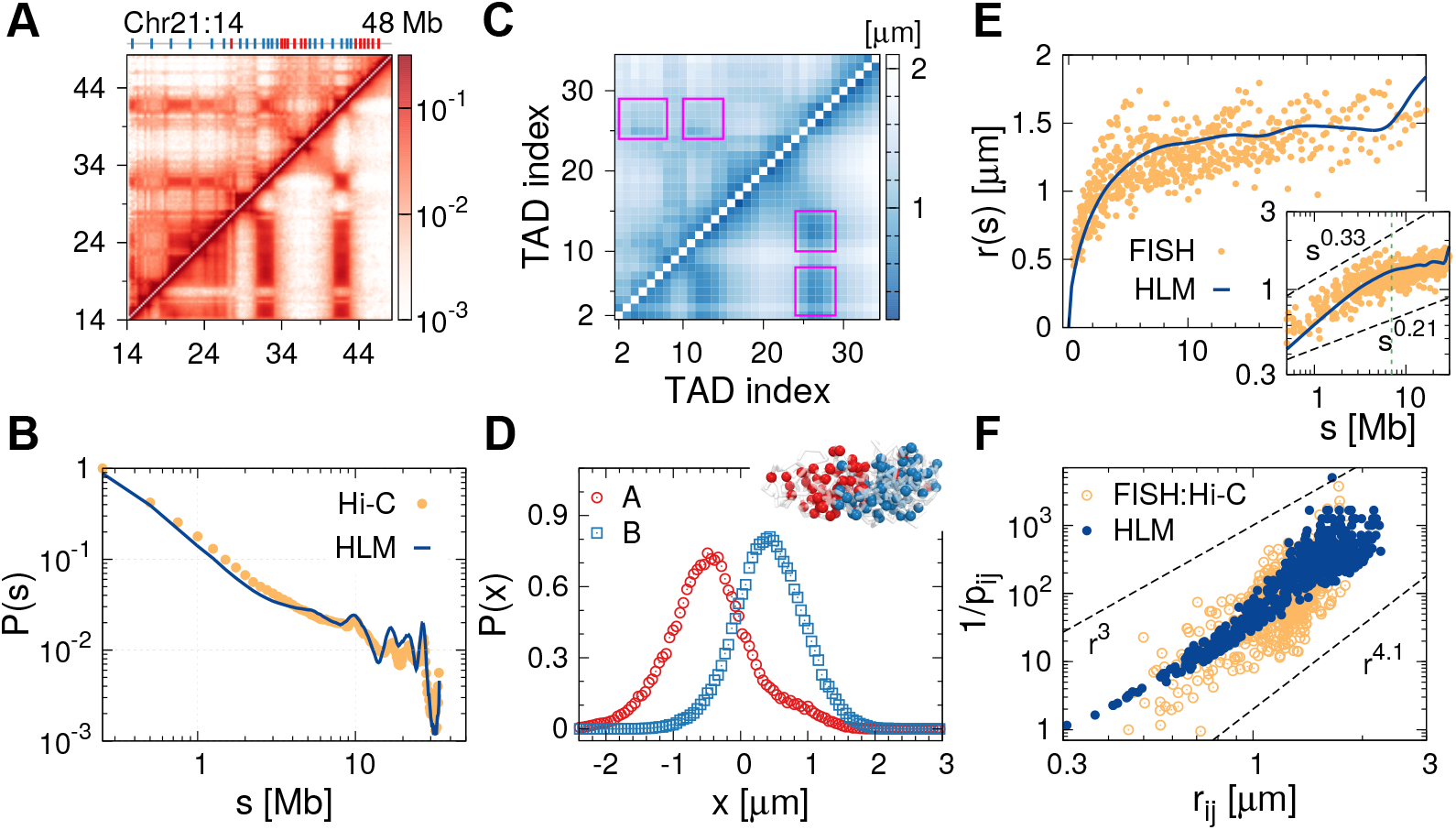
A 34 Mb-genomic region of chr21 in IMR90 cells modeled by HLM. (A) Heatmap of contact probabilities from Hi-C (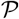, upper diagonal part) and HLM (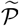, lower diagonal part). The Pearson correlation between 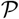 and 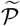 is 0.97. The positions of TADs are displayed above the heatmap, labeled by sticks. The type of each domain, A and B, is depicted in red and blue. (B) Mean contact probability *P*(*s*) calculated from Hi-C (orange data) and HLM (blue line). (C) The heatmap of inter-TAD distances measured by FISH (upper diagonal part) is compared with that calculated from HLM (lower diagonal part). (D) Distributions of A- and B-type TADs projected on the *x*-axis, along which the geometric centers of different types of TADs are aligned, are indicative of the microphase separation. An ensemble of structures are also shown. (E) Intrachain end-to-end distance *r*(*s*) as a function of arc-length *s* from FISH (orange data) and HLM (blue line). The inset shows *r*(*s*) in log-log scale. (F) Inverse of pairwise contact probability between TADs, 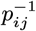, versus inter-TAD distance *r_ij_* in log-log scale.

The heatmap calculated for inter-TAD distances using the HLM-generated conformational ensemble (lower diagonal part of Fig. 1C) can directly be compared with the FISH measurement (upper diagonal part). The square block pattern along the diagonal axis of the heatmap indicates that 4–5 adjacent TADs constitute an aggregate, reminiscent of meta-TAD^30^, and the patterns in the off-diagonal part (highlighted by the magenta boxes) suggest long range clustering of TADs. The error of the inter-TAD distance heatmap relative to FISH is 0.184, which is comparable to the value of GEM model^38^ and better than others (see Fig. 4D in Ref.^38^). A principal component analysis of this matrix (top left part of the matrix in Fig. 1C) divides TADs into A/B types^23^. Aligning the geometric centers of HLM-generated A- and B-type TADs parallel to the *x*-axis highlights a polarized organization of A- and B-type TADs (see Fig. 1D)^23^.

The intrachain end-to-end distance 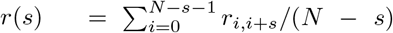 displays a scale-dependent scaling relationship with the genomic distance *s, r*(*s*) ~ *s^ν^* (Fig. 1E). In qualitative agreement with the FISH measurement^23^, there is a crossover around *s* = 7 Mb, such that *ν* ≈ 1/3 for *s* < 7 Mb and *ν* ≈ 0.21 for *s* > 7 Mb.

We explore the relationship between contact probability *p_ij_* and the corresponding distance *r_ij_* of two loci. It is expected that the looping probability of polymer is inversely proportional to the volume of space (*V*) explored by the two loci as *P*_loop_ ~ 1/*V*. Since the volume *V* scales with the spatial separation (*R*) between the two loci in *d*-dimension as *V* ~ *R^d^*, it follows that^54–56^

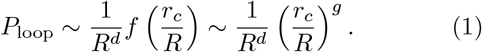

The correlation hole exponent *g* is *g* = 0 for a Gaussian chain^57^. According to the Flory theorem^58–61^, the ideal chain statistics is a good approximation for a chain in polymer melts or for a subchain in a fully equilibrated globule. Since *d* = 3 for 3D, we expect *P*_loop_ ~ *R*^−3^, or equivalently 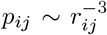 (see also Fig. S1B). In fact, this scaling relation is observed for the data point generated by HLM for *r_ij_* < 1 *μ*m (Fig. 1F). Although Wang *et al*., who combined Hi-C and FISH data, reported a scaling relation of 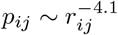 for the entire range, it is not clear whether the relation can straightforwardly be extended to the range of *r_ij_* < 1 *μ*m where the data point from their measurement might be less accurate. According to the HLM-generated data a more proper scaling should be 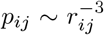 for *r_ij_* < 1 *μ*m and 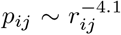 for *r_ij_* > 1 *μ*m.

Next, to demonstrate another analysis on FISH measurement, we applied HLM to the q-arm of chr11 in IMR90 cells, whose intrachain pairwise distances between genomic loci had been measured with FISH^28, 64^ (see Table S2 for the position of FISH probes in the genome and in the model). The model produces the contact probability matrix 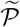 with a Pearson correlation (PC) of 0.98 relative to Hi-C data 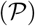 (see Figs. S3A, S3B, and SI for discussion of PC in comparison to other alternative method). HLM enables us to calculate the spatial distances between specific pairs of loci (Fig. S3C), with a mean relative error of 0.189 (with respect to FISH data). The HLM-generated structural ensemble also indicates that compared to the gene-poor and transcriptionally inactive anti-ridge domain, the transcriptionally active ridge domain is less compact, less spherical, and has a rougher domain surface (Figs. S3D-F), all of which are in agreement with the FISH experiment^64^. Modeling another 30 Mb region on chr1 of IMR90 cells leads to similar results (Fig. S4 and Table S3).

### Visualization of chromatin globules

#### *α*-globin gene

*Cis*-regulatory elements generally mediate the transcription of neighboring genes within a range smaller than 1 Mb^65^. The *α*-globin gene domain, a 500 kb-genomic region known as ENm008 located at the left telomere of human chr16, has previously been studied to decipher the relationship between chromatin structure and transcription activity^37, 52, 53^. RNA-seq data^62, 66, 67^ indicate that the *α*-globin genes (including *ζ*-, *μ*-, *α*2-, *α*1- and *θ*-globin genes) are expressed in K562 cell lines, but silenced in GM12878 (tracks on the left side of the Hi-C heatmaps in Fig. 2A). According to 3C/5C measurements^52, 68^, the *α*-globin gene forms long-range looping interactions with multiple regulatory elements upon gene activation. Among them, of particular interest is one of the DNase I-hypersensitive sites (DHS), HS40, located at ~ 70 kb upstream of the *α*1 gene.

**FIG. 2.**
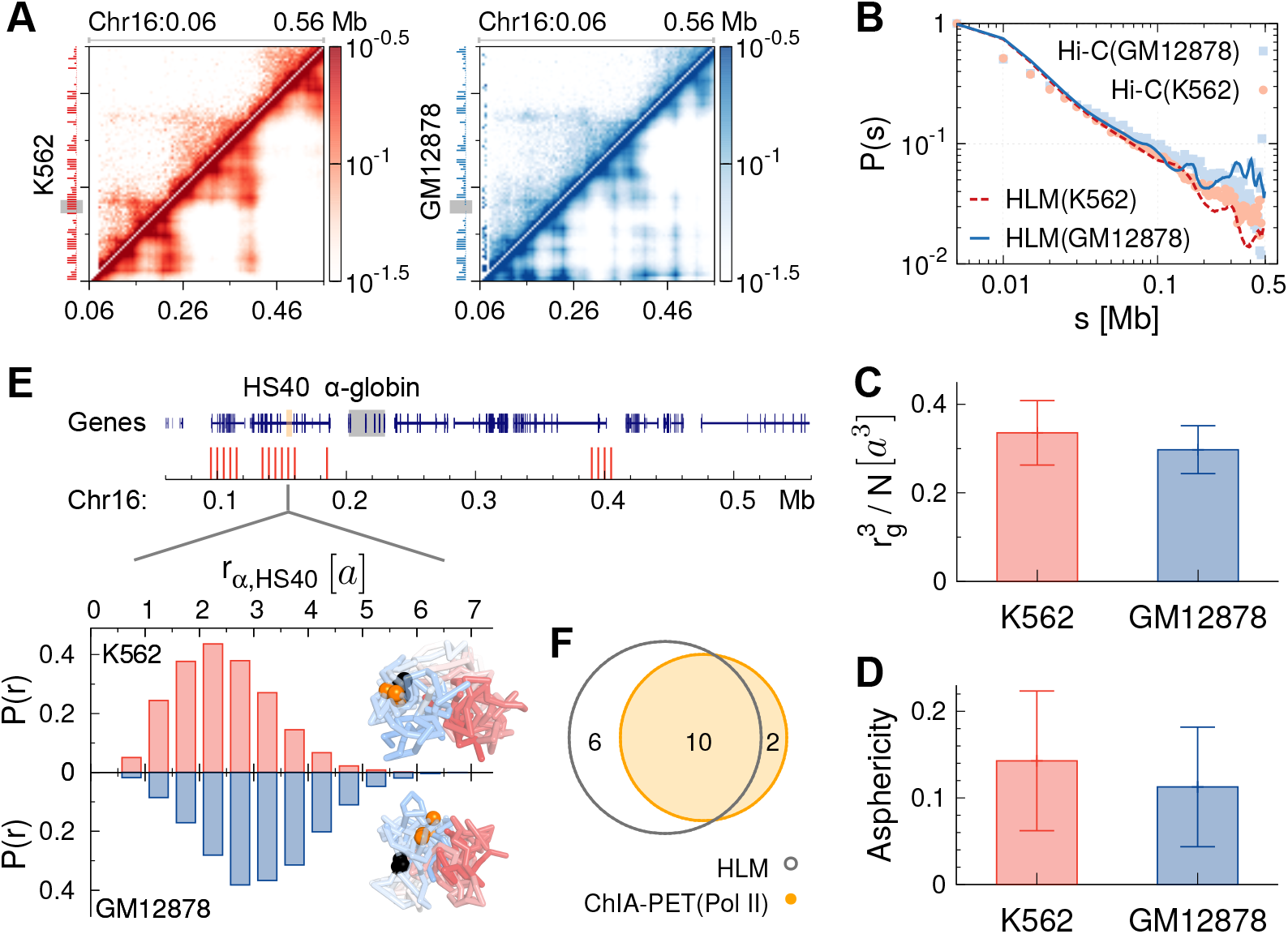
*α*-globin gene domain modeled by HLM for two different cell lines. (A) Heatmap 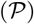 of contact probabilities measured by Hi-C (upper diagonal part) and the corresponding map 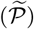 obtained from HLM (lower diagonal part), in K562 (left) and GM12878 (right) cells. RNA-Seq signals^62^ is displayed on the left side of the heatmaps, and the location of *α*-globin gene is depicted in gray shade. (B) Mean contact probability *P*(*s*). (C) Compactness, and (D) asphericity of the domain. (E) Gemonic positions of the loci, closer to the *α*1 gene in K562 than GM12878 cells, are labeled using red sticks. Contrasted below are the distance distributions between *α*1 gene and HS40, *P*(*r*_*q*, HS40_), for two cell lines. For each cell line, an ensemble of structures are shown for comparison with chains colored by the genomic position from the telomere (blue) to centromere (red). The *α*-globin gene and HS40 are rendered as a black and orange sphere, respectively. (F) Pol II-mediated chromatin interactions^50^, which involves *α*-globin genes and specific to K562 cells, are compared with the model.

The HLM-generated structural ensemble at 5 kb resolution for ENm008 of two cell lines (K562 and GM12878) suggests that the contact probability *P*(*s*) decreases slightly faster in K562 than in GM12878 cells at large s (Fig. 2B). The *α*-globin domains of K562 and GM12878 cell lines visualized with FISH^52^ indicates that K562 is less compact than GM12878, which is confirmed straightforwardly by the compactness calculated using the HLM-generated structures (Fig. 2C). Compared with GM12878 cells, the *α*-globin domain in K562 cells adopt a less spherical shape (Fig. 2D)^52, 53^.

Next, we examined the changes in the distances between the *α*1-globin gene and other loci upon activation of the gene. Even though the whole domain in K562 cells is relatively more expanded, HS40 is closer to the *α*1 gene in K562 than in GM12878 cells (Fig. 2E), which is consistent with the expectation based on the higher contact enrichment between HS40 and *α*1 gene observed in K562 by 3C/5C measurements (e.g., Fig. 2 in Ref.^52^). Through inter-cell line comparison between K562 and GM12878 for the rest of the region using distance distribution to the *α*-globin gene locus, we identified a group of loci other than HS40 that are significantly closer to *α*-globin genes in K562 cells (Mann-Whitney U test, *p* < 1 × 10^−5^). Their genomic positions are marked using red sticks in Fig. 2E. According to the independent ChIA-PET experiments^50, 51^ designed to capture the chromatin loop interactions mediated by specific protein factors, the structural variation associated with *α*-globin genes is mainly orchestrated by Pol II (see Table S4). HLM captures 83% of Pol II-mediated chromatin loops specific to K562 cells (Fig. 2F).

Taken together, HLM captures both the tissue-specific variation in the global packing of the *α*-globin gene domain, and variation in the structure of gene locus. The multiple K562-specific interactions, substantiated by HLM, suggest that a cooperative action of multiple regulatory elements including HS40 is responsible for the activation of *α*-globin genes^37^. HLM-generated conformations indeed confirm the notion of chromatin globule proposed in Ref.^52^.

#### SOX2 gene

As an another example of transcription-dependent chromatin folding, we studied the human SOX2 gene locus which encodes a transcription factor involving the regulation of embryonic development. The SOX2 gene is transcribed in human embryonic stem cells (hESCs), but not in umbilical vein epithelial cells (HUVECs) (Fig. 3A). To compare the results from HLM with a recent modeling study^32^, we measured the distances between SOX2 gene and two possible regulatory elements located at regions ~800 kb upstream (US) and ~650 kb downstream (DS). Whereas both elements are closer to the SOX2 locus in transcriptionally active hESCs than in inactive HUVECs, the chromatin fiber is less compact in hESCs (Fig. 3D, see also the snapshots in Figs. 3E and F). HLM-generated structures demonstrate the dependence of chromatin folding on the transcription level at SOX2 gene loci, and this trend comports well with the prediction made in Ref.^32^ that also employed polymer model simulation.

**FIG. 3.**
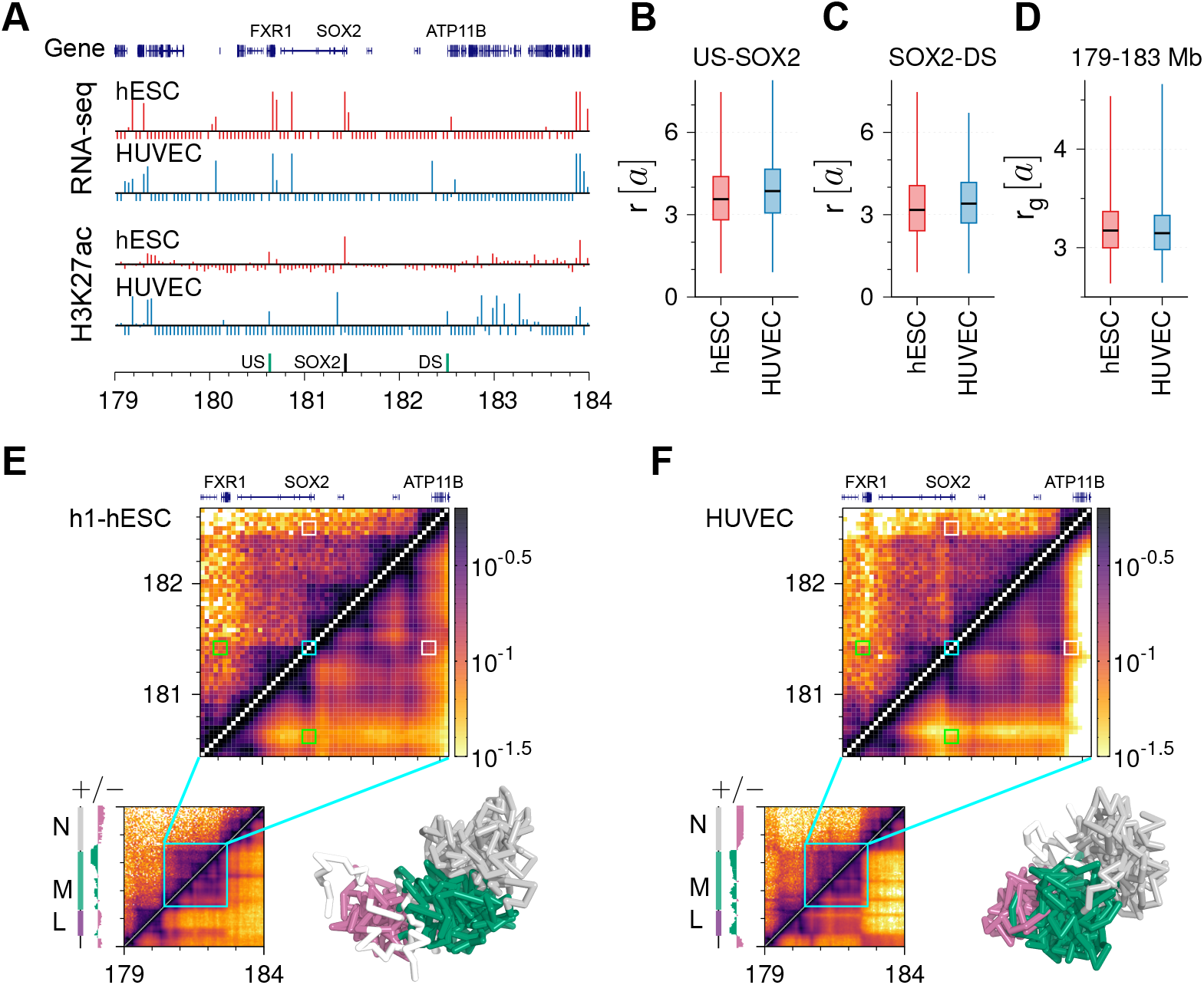
Comparison of a 5 Mb genomic region on chr3 modeled by HLM between hESCs and HUVECs, which includes SOX2 gene. (A) Genes annotated in this region, aligned with RNA-seq^62^ and H3K27ac signals^63^ of two cell lines. The genomic positions of three “simulated” FISH probes^32^ are labeled in the bottom track. (B) The distance between US and SOX2, (C) distance between SOX2 and DS, and (D) the gyration radius calculated from our model. (E) Heatmap of contact probabilities for hESCs measured by Hi-C^47^ (upper diagonal part) and calculated from HLM (lower diagonal part). Based on the first principal component of the significance matrix (track on the left side of heatmap), we divided the region into three domains, and colored the chromatin chain accordingly in the snapshot of a typical structural ensemble. (F) Analysis was carried out for HUVECs with Hi-C data from Ref.^3^.

### Chromatin interactions at complex genomic loci

The efficacy of HLM was further tested for the genomic loci of Pax6 gene that involve the development of mouse neural tissues. Flanked by two neighboring genes (Pax6os1 and Elp4), the expression level of Pax6 gene is considered to be regulated by multiple long-range elements, including two regulatory regions located at ~50 kb upstream (URR) and ~95 kb downstream (DRR) (Fig. 4A). The DRR contains several DNase I-hypersensitive sites and the SIMO enhancer, which was identified in transgenic reporter gene studies of developing mouse embryos^71, 72^. Another cis-regulatory element PE3 within URR has recently been identified from mouse pancreatic *β*-cells (*β*-TC3)^70^.

**FIG. 4.**
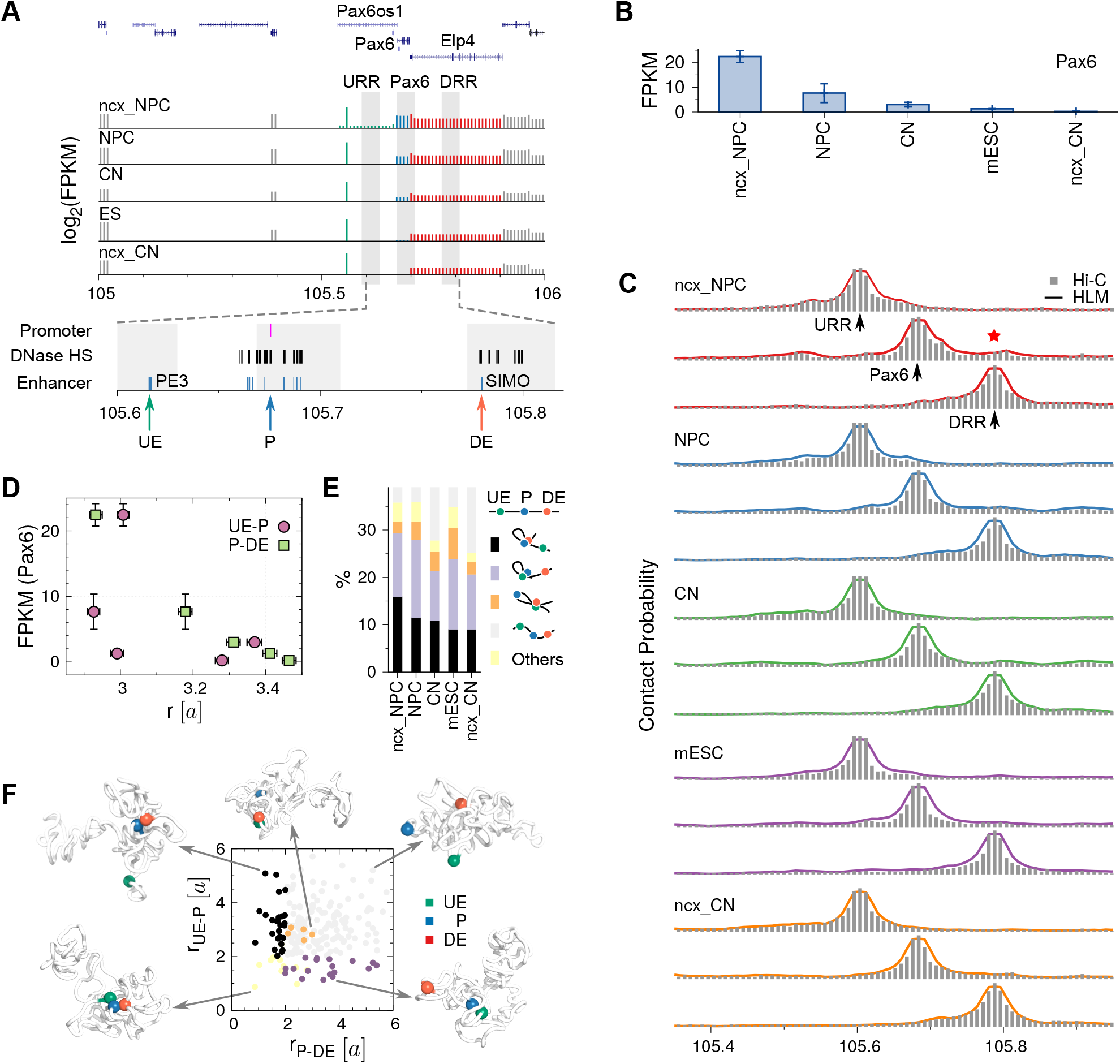
Pax6 gene locus modeled by HLM for five different mouse cell types. (A) Genes in a 1 Mb region on chr2 where the genomic position of URR, Pax6, and DRR are labeled with gray shade, in alignment with the FPKM score measured from RNA-seq analysis^48^. Pax6 gene promoters, enhancers and nearby DNase I hypersensitive sites are zoomed in at the bottom^69, 70^, where the positions of the upstream enhancer (UE), promoter (P) and downstream enhancer (DE) are marked with arrows. (B) Expression levels of the Pax6 gene in different cell types. (C) Contact profiles for five different cell types. The profiles were constructed using Hi-C data (gray bars) for URR, Pax6, and DRR (from top to bottom) relative to other genomic regions and calculated using HLM (solid lines). (D) Pax6 expression level as a function of the average distance between two enhancers (UE and DE) and the promoter (P). (E) Percentage of chromatin conformations belonging to each group classified based on the distances betwen UE, P, and DE. (F) Scatter plot of the distances *r*_P-DE_ and *r*_UE-P_ of 200 structures, which were randomly selected from the conformational ensemble of ncx_NPC cells. Typical structures are presented for each group where the three sites are labeled using different colors.

A study combining Capture-C, FISH and simulations^32^ has reported a non-trivial correlation between the expression level of Pax6 gene and the spatial separation from Pax6 gene to URR and DRR. Among the three types of mouse cells (*β*-TC3, MV+ and RAG cells) studied in Ref.^32^, Pax6 gene maintained the largest separation from DRR in the *β*-TC3 cells that displayed the highest expression level of Pax6. Therefore, it was suggested^32^ that the enhancer at DRR is not involved in upregulation of Pax6 in *β*-TC3 cells, or that some unclear upregulation mechanisms that do not require the spatial proximity to enhancers are responsible for the activity of Pax6 gene.

To study the origin of complex interplay between Pax6 gene and neighboring genetic elements, we applied HLM to the same genomic region of five different mouse cell types whose Hi-C data are currently available: (i) embryonic stem cells (mESCs), (ii) neural progenitors (NPCs), (iii) cortical neurons (CNs), (iv) ncx_NPC, and (v) ncx_CN, where the prefix “ncx_ “ indicates that the cells are directly purified from the developing mouse embryonic neocortex *in vivo*. Each cell type displays distinct transcriptional activity patterns of Pax6 and its neighboring genes^48^ (Fig. 4A). According to the FPKM scores from RNA-seq analysis (Fig. 4 B), the five cell types display Pax6 activity in the following order: ncx_NPC > NPC > CN > ES > ncx_CN.

The contact probabilities calculated from our HLM-generated conformations reasonably reproduce the Hi-C data at a resolution of 8 kb^48^ (see Table I and Fig. S5). The Hi-C contact profiles of three genomic loci (URR, Pax6, and DRR) with other genomic regions (histograms in Fig. 4C) are well captured by HLM-generated conformations (lines in Fig. 4C). Compared with the distance of Pax gene promoter (P) to the upstream enhancer (UE), Pax6 gene activity is better correlated with the distance to the downstream enhancer (DE) (see Fig. 4D); the closer to the DE, the higher the Pax gene activity is. The highest Pax gene activity is seen in ncx_NPC. Notice that the most enriched Hi-C contacts between Pax6 and DRR is indeed found in ncx_NPC, which is marked with a red star in Fig. 4 C. We note that our finding on contacts between Pax6 and DRR is in contrast to that based on *β*-TC3 cells (see Fig. 2 A in Ref.^32^). This however underscores that the mechanism or the chromatin conformations responsible for the Pax6 gene activity depends strongly on the cell-type: At least the mechanism of Pax6 gene regulation in ncx_NPC cells differs clearly from that in *β*-TC3 cells.

Next, given that Hi-C data is obtained from a collection of millions of cells, heterogeneity of chromatin conformations is inevitable in analyses, which has indeed been highlighted in Ref.^32^. To characterize the heterogeneity in the HLM-generated conformational ensembles, we classified each chromatin structure into five groups based on the separations between the Pax6 gene promoter (P) and two enhancers (UE and DE) (Fig. 4E). To visualize the conformational diversity, we randomly selected 200 structures and characterized by the promoter-enhancer distances (Fig. 4 F). Except for the “gray” group where all three separations are large, the population of conformational ensemble consists mainly of the “black” group (P is close to DE but not to UE), and the “purple” group (P is close to UE but not to DE) which are suspected to be responsible for high expression level of Pax6 gene. In consistent with our analysis on the ensemble-averaged distance to enhancers for different cells (Fig. 4 D), the proportion of “black” group shows a decreasing trend as Pax6 becomes less active (Fig. 4 E), suggesting a more important role of DE than UE in regulating Pax6 gene for the five cell lines.

While an indirect upregulation of Pax6 gene by DRR as seen in *β*-TC3 cells^32^ cannot entirely be ruled out, the correlation of gene activity level with the spatial proximity of Pax6 gene to DRR is clearly demonstrated, at least, across the five cell lines that we studied using HLM. The mechanism of indirect upregulation and the mechanism of cell type-dependent choices deserve further study.

### Chromosome in different phases of cell cycle

Most Hi-C data are obtained over a population of ‘unphased’ cells. Here, we employ HLM to model the global architecture of chromosome at different phases of cell cycle during the interphase, based on single-cell Hi-C^4^. Accumulating the data from tens to hundreds of binary contact matrices of single cells into an input matrix 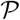, we built 500 kb-resolution model of chromosome for the post-M, early-S, mid-S, late-S/G2, and pre-M phases of chr19 in mESC (above the diagonal in Fig. 5A). 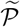 matrices computed using HLM (below the diagonal in Fig. 5A) display reasonable correlation with the original Hi-C data (Pearson correlation, PC > 0.9) except for the post-M phase (PC = 0.77); unlike other phases, the lower PC value with the 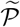-matrix at the post-M phase, characterized with uniform and featureless pattern, is due to the smaller number of sampling cells (*N_c_*).

**FIG. 5.**
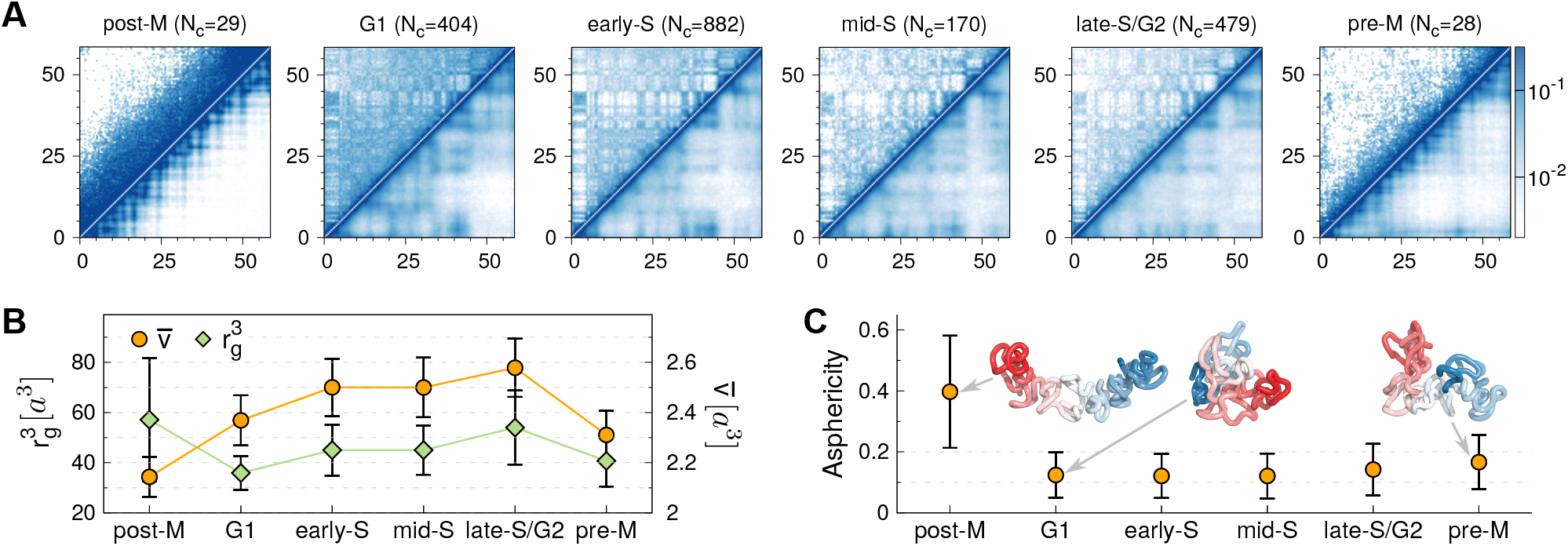
Chr19 of mESC modeled by HLM at 500 kb resolution. (A) Heatmap of contact probabilities from Hi-C (upper diagonal part) and HLM (lower diagonal part). From post-M phase to pre-M phase, Pearson correlations (PCs) are 0.77, 0.96, 0.96, 0.96, 0.97 and 0.91, respectively. The Hi-C matrices are the outcomes from a sum of *N_c_* binary contact matrices of single cells in the same phases of the cell cycle. (B) 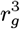 and the average volume 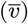 occupied by a single monomer. (C) Asphericity of the chromosome in different phases along the cell cycle. Depicted at post-M, G1 and pre-M phases are the snapshots of the HLM-generated structure which are colored from the centromere (blue) to telomore (red).

The local compactness of the chromosome conformation was quantified in terms of the average volume occupied by a single monomer 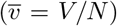 based on the Voronoi tessellation (Fig. 5B). After the mitosis, the chromosome continues to expand until the late-S/G2 phase. The gyration radius also captures this trend (Fig. 5B), except that the model has the largest value of *r_g_* in post-M phase. A partial condensation of the chain (decreases in 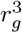 and 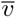) is observed before entering the pre-M phase. This decondensation-condensation cycle is also captured with the asphericity of structures generated from HLM (Fig. 5C), which decreases dramatically from the post-M to G1 phases and then increases gradually after the G1 phase. The same conclusion can be drawn from the probability density of pairwise distance between monomers (see Fig. S6).

## DISCUSSION

HLM is similar to previous polymer models of chromatin, which also convert information of spatial proximity into harmonic restraints between monomers^25, 73, 74^. In order to demonstrate that the choice of energy potential in HLM is optimal over other alternatives, we examined HLM and its three variants on a 10 Mb genomic region on chr5 of GM12878 cells (Fig. S7). Unlike the HLM which faithfully reproduced the domain edges of enriched contacts observed by Hi-C (highlighted by cyan boxes in Fig. S7A), which was regarded as a distinct feature of loop extrusion^14^, two alternative copolymer models, which retain uniform strength of loop interaction, could not properly reproduce the diagonal-block patterns of Hi-C data (Fig. S7B and C). In a homopolymer model, where *χ*_−,−_, *χ*_−,+_, and *χ*_−,+_ are all set to 1 (see Methods), the long-range checkboard pattern was not reproduced (Fig. S7D). The Pearson correlation of contact probabilities contrasted between Hi-C and other models at different genomic separations shows that HLM outperforms others (Fig. S7E).

As shown for different chromosomes, cell types, species with a flexible choice of model resolution, one of the greatest advantages of HLM is its versatile application. While all of the output conformations exhibit great variability (see discussions in SI, Fig. S8, and Fig. 4F), the population-sampled contact map faithfully reproduces the input Hi-C data. For a given Hi-C data, the two sets of model parameters 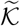 and {*χ_t_i_, t_j__*} can be determined in a few minutes using a personal computer without any manual intervention (Table I).

In summary, we demonstrated that HLM is a computationally efficient approach with which to investigate the genome function. The conformational ensemble generated by HLM shows that depending on the chromatin states, different types of chromatin domains have different compactness and shapes, and spatial phase separation between domains takes places in human genome. The inter-cell line comparison of human *α*-globin and SOX2 loci shows that while the sub-megabase gene domain becomes less compact upon gene activation, the most critical regulatory element comes closer to the gene, and that its expression is likely affected by many other elements. The activity of Pax6 gene in a complex genetic environment is mostly modulated by the proximity between Pax6 promoter and the downstream enhancer, while the distance to the upstream regulatory element shows non-monotonic variations with its activity for the cell types we studies. HLM was also used to visualize the cell cycle dynamics of chromosome organization based on single-cell Hi-C. Although HLM is not designed based on assumptions of molecular mechanisms of genome organization, the principle of transcription regulation can be inferred from the changes of chromatin conformations. With Hi-C data being accumulated, HLM would be of great use to provide complementary structural information, which are not easily accessible to current experiments.

## METHODS

### Description of HLM

The full energy potential of HLM consists of two parts.
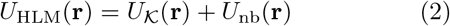

In what follows, we delineate the first and second terms of Eq. 2 (see SI for technical details).

First, decomposed into two parts, 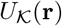 describes the harmonic constraints on a chain of *N* monomers^27^,

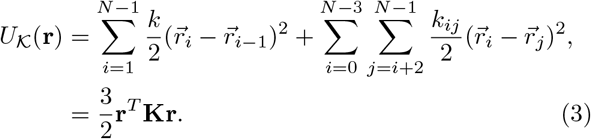

where successive monomers along the backbone and non-successive monomers forming loops are both harmonically restrained. In the second line, 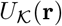 is written in a compact form with 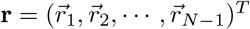 and **K** representing the Kirchhoff matrix. **K** can be built from the interaction strength matrix 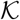 that takes 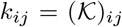 as its matrix element. The interaction strengths ought to be non-negative (*k_ij_* ≥ 0) for all *i* and *j*-th monomer pairs. In HLM, if *k_ij_* ≠ 0 then the *i* and *j*-th monomer has a potential to form a (chromatin) loop. After removing the translational degrees of freedom by setting 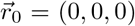 on Eq. 3, we obtain the probability density of pairwise distance as^27^

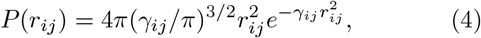

where

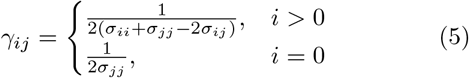

and 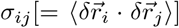 is the covariance between the positions of *i* and *j*-th monomers, which can be obtained from an inverse of **K**-matrix as

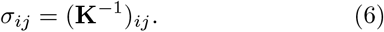

One can obtain the contact probability *p_ij_* by integrating the pairwise distance *P*(*r_ij_*) (Eq. 4) up to a certain capture radius (*r_c_*)^75, 76^, 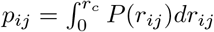, which gives

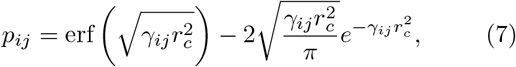

where 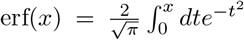. Therefore, a one-to-one analytical mapping between *p_ij_* and *k_ij_* follows from the precise mappings between *p_ij_* and *σ_ij_* from Eqs 7 and 5, and between *σ_ij_* and *k_ij_* from Eq. 6.

Although it is tempting to directly use the mathematical relation between *p_ij_* and *k_ij_* to obtain 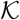 from Hi-C data, there is an unavoidable numerical issue (see SI Text and Figs. S9–S11 for details). In practice, we calculate 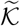-matrix that approximates 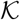 by selecting only the *significant* contacts in 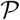. More specifically, we evaluate the significance of contact probability *p_ij_* by calculating *z_ij_*, which is defined as (see the matrix elements in the upper diagonal part of Fig. 6B):

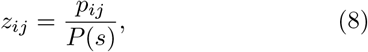

where 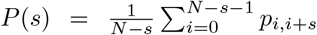 is the mean contact probability for monomer pairs separated by the arc length *s* along the contour. The greater the value of *z_ij_*, the contacts are deemed more *significant*. We then select top 2*N* (*i, j*) pairs ranked in terms of the values of *z_ij_* (> 1) (the matrix elements in the lower diagonal part of Fig. 6B). For these 2*N* pairs whose contact probability *p_ij_* is given in 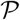, the precise value of 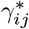 (or equivalently 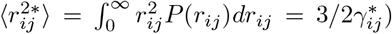 can be determined using Eq. 7. Then, starting from a Rouse chain configuration as an initial input, we add non-successive bonds with varying interaction strengths (0 ≤ *k_ij_* ≤ 10 *k_B_T*/*a*^2^) until we minimize the objective function 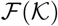

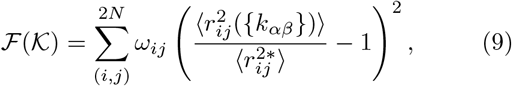

so as to determine the optimal values of 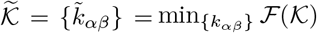. Here the weight factor *ω*_ij_, which is used to normalize the statistical bias from chromatin loops of different sizes, is defined as

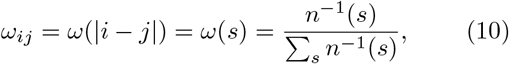

where 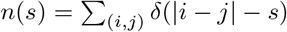 is the number of loops of size *s*. The gradient-descent algorithm (L-BFGS-B method in SciPy package) was used to determine the optimal parameters 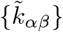. A fully convergent solution of 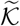-matrix (Fig. 6C) could be obtained within *a few minutes* when *N* was not too large (≤ 200). This 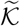-matrix determining process, termed *constrained optimization*, faithfully reproduces the original 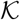 matrix with a relative error smaller than 5% (see also Figs. S10-S12).

**FIG. 6.**
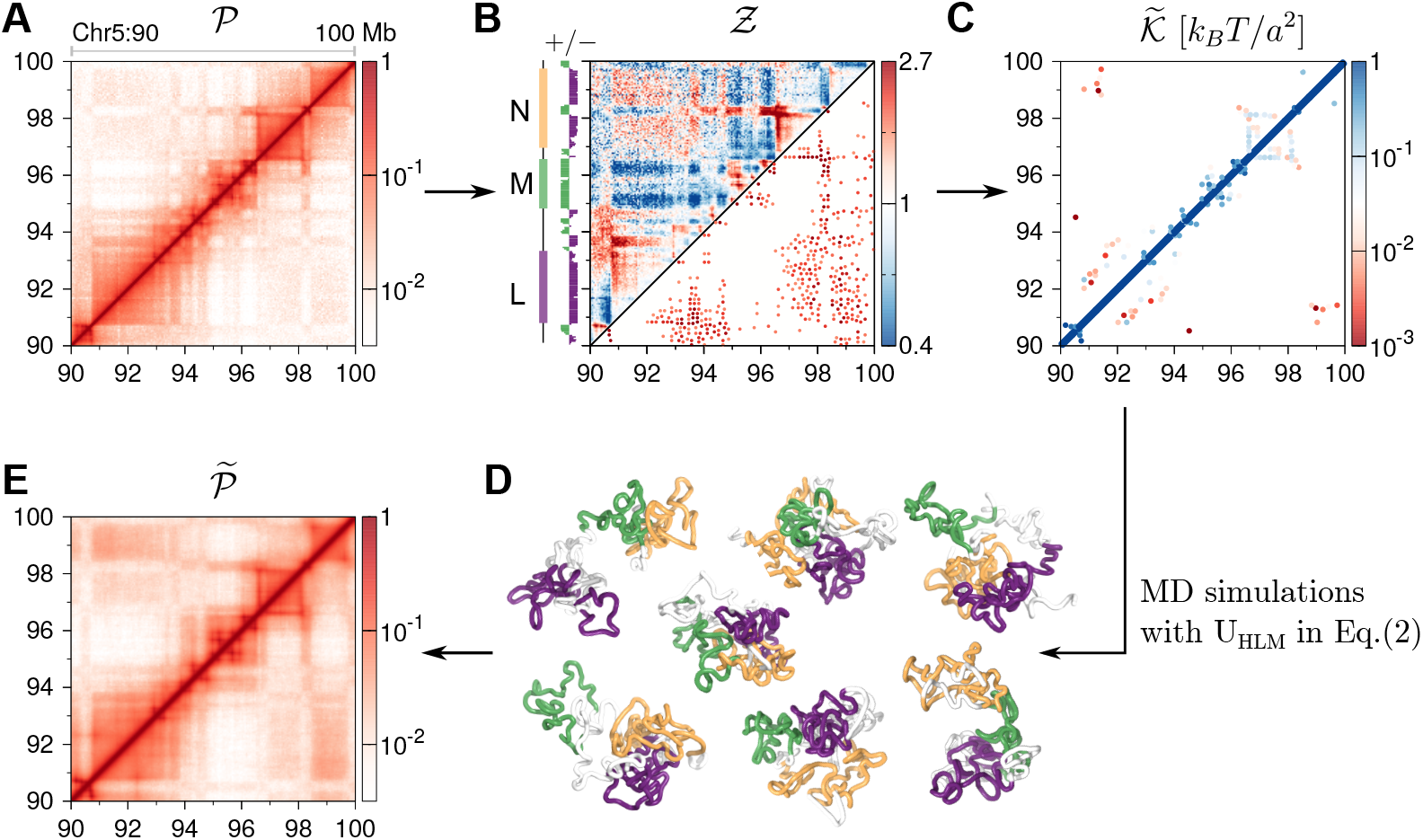
The pipeline of HLM. (A) Contact probability matrix 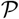 of a 10 Mb-genomic region of chr5 in GM12878 cells from Hi-C. (B) 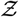 matrix (above the diagonal) calculated from Eq. 8. The significant contacts selected from 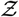 (below the diagonal). The sign of the first principal component of 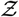 is provided on the left hand side of the panel, and we divide the whole chromosome domain into “L” (purple),”M” (green) and “N” (orange), accordingly. (C) Interaction strength matrix 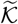 calculated by the constrained optimization. (D) Conformational ensemble of chromosomes generated from HLM potential defined by 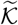 and 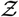 (Eq. 2). Illustrated are the chromosome conformations with L, M, N domains colored in purple, green, orange, respectively, following the domain labels assigned in Fig. 6B. (E) 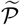-matrix calculated using a conformational ensemble produced from MD simulations. Pearson correlation between 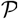 (Fig. 6A) and 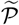 (Fig. 6E) is 0.96.

After obtaining 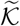 (Fig. 6C), and hence 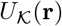, we added a non-bonded interaction term *U*_nb_(r), defined for all *i* and *j* pairs to the full energy potential *U*_HLM_(*r*) (Eq.2):

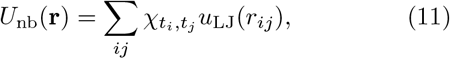

where *u*_LJ_(*r*) is the Lennard-Jones potential truncated for *r* ≥ *r_c_* where *r_c_* = 5*a*/2 with ∊ = 0.45 *k_B_T*,

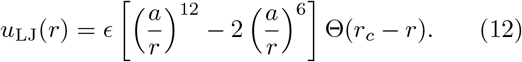

If ∊ = *∊_θ_*(= 0.34 *k_B_T*) with *χ_t_i_, t_j__* = 1, then *U*_nb_(r) leads to *θ*-solvent condition for infinitely long chain, putting the second virial coefficient to zero, i.e., 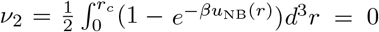. We chose ∊(= 0.45 *k_B_T*) slightly greater than *∊_θ_* and assigned loci-pair-type-dependent prefactor *χ_t_i_, t_j__*. Each monomer *i* is assigned with a type *t*, either “−” or “+”, based on the sign of the first principal component of 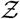 (see the track on top of Fig. 6B). The value of prefactor *χ_t_i_, t_j__* (> 0), depending on the types of two loci *i* and *j* which are either *t_i_t_j_* = ++, −−, or −+, are evaluated by averaging over all the monomer pairs of the corresponding types, such that *χ_p, q_* = 〈*z_ij_*〉_*t_i_*=*p, t_j_*=*q*_, The values of *χ_t_i_, t_j__* are determined based on a given Hi-C data. For the case shown in Fig. 6, we obtain *χ*_−,−_ = 1.18, *χ*_−,+_ = 0.79, and χ+,+ = 1.19. According to the Flory-Huggins theory^57^, the condition 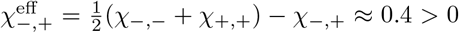 leads to microphase separation between + and − type loci, which indeed is realized and reflected in the characteristic checkerboard pattern of Hi-C data. It should be noted, however, that the classification of type −/+ is not necessarily identical to the A/B compartment of chromatin. Whereas A/B compartments are genome-wide characteristics usually defined based on Hi-C data at low (Mb) resolutions^2, 3^, the monomers in HLM can be always classified into types −/+ regardless the resolution of the model.

Finally, we sampled 3D chromosome structures using molecular dynamics simulation implementing the full energy potential *U*_HLM_(*r*) and calculated the contact probability matrix based on HLM-generated conformational ensemble. In the specific example demonstrated for the Hi-C data of 10 Mb-genomic region of chr5 in GM12878 cell line (Fig. 6), 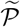 (Fig. 6E) obtained from HLM-generated chromosome conformations (Fig. 6D, see also the clustering analysis which highlights the conformational variability of chromosomes in SI text and Fig. S8) displays a notable resemblance to the input 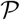 (Fig. 6A) (Pearson correlation of 0.96; Spearman correlation of 0.92). Despite the simplicity of HLM potential (Eq. 2), the similarity between 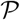 and 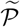, as well as the chromosome conformations ensemble generated during the procedure is remarkable.

### Structure characterization

We quantified the structural feature of HLM-generated chromosome ensemble, by means of several quantities:

i. The compactness of a (sub-)chain of length *N* is quantified in terms of 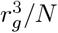, where *r_g_* is the gyration radius of the (sub-)chain.
ii. The asphericity (*A*) is calculated by 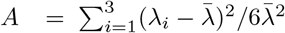 where *λ_i_*(*i* = 1, 2, 3) are the three eigenvalues of the moment of inertia tensor, and 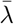 is their mean^77, 78^. *A* = 0 for a sphere, and *A* > 0 for a non-spherical shape.
iii. The roughness of the surface of a (sub-)chain, was evaluated using the Voronoi diagram^79^ that tessellates the 3D space occupied by the chain. A upper bound for the volume of each monomer was set using a dodecahedron with a diameter of 2*a*, The Voronoi diagram provides a well-defined volume *V* and surface area *S* of the (sub-)chain. Since the surface area of a perfect sphere with the volume *V* is *S*_0_ = (36*πV*^2^)^1/3^, we quantified the surface roughness using *S*/*S*_0_ ≥ 1.
iv. To visualize an ensemble of structures with considerable variability, we first divided the chain into a few segments (domains). Next, the distribution of the distances between the geometric centers of these domains were computed based on the ensemble of structures. Several configurations of chromosomes were then randomly selected from the most populated state (in terms of interdomain distances), aligned, and rendered.

## ACKNOWLEDGMENTS

We thank the Center for Advanced Computation in KIAS for providing computing resources. CH acknowledges a partial support from the National Research Foundation of Korea (NRF-2018R1A2B3001690).

## References

1 J. Dekker, K. Rippe, M. Dekker, and N. Kleckner, “Capturing chromosome conformation,” Science 295, 1306–1311 (2002).

2 E. Lieberman-Aiden, N. L. van Berkum, L. Williams, M. Imakaev, T. Ragoczy, A. Telling, I. Amit, B. R. Lajoie, P. J. Sabo, M. O. Dorschner, R. Sandstrom, B. Bernstein, M. A. Bender, M. Groudine, A. Gnirke, J. Stamatoyannopoulos, L. A. Mirny, E. S. Lander, and J. Dekker, “Comprehensive mapping of long-range interactions reveals folding principles of the human genome,” Science 326, 289–293 (2009).

3 S. S. P. Rao, M. H. Huntley, N. C. Durand, E. K. Stamenova, I. D. Bochkov, J. T. Robinson, A. L. Sanborn, I. Machol, A. D. Omer, E. S. Lander, and E. L. Aiden, “A 3D map of the human genome at kilobase resolution reveals principles of chromatin looping,” Cell 159, 1665–1680 (2014).

4 T. Nagano, Y. Lubling, C. Várnai, C. Dudley, W. Leung, Y. Baran, N. Mendelson Cohen, S. Wingett, P. Fraser, and A. Tanay, “Cell-cycle dynamics of chromosomal organization at single-cell resolution,” Nature 547, 61–67 (2017).

5 Z. Du, H. Zheng, B. Huang, R. Ma, J. Wu, X. Zhang, J. He, Y. Xiang, Q. Wang, Y. Li, J. Ma, X. Zhang, K. Zhang, Y. Wang, M. Q. Zhang, J. Gao, J. R. Dixon, X. Wang, J. Zeng, and W. Xie, “Allelic reprogramming of 3D chromatin architecture during early mammalian development,” Nature 547, 232–235 (2017).

6 C. M. Rivera and B. Ren, “Mapping human epigenomes,” Cell 155, 39–55 (2013).

7 F. Jin, Y. Li, J. R. Dixon, S. Selvaraj, Z. Ye, A. Y. Lee, C.-A. Yen, A. D. Schmitt, C. A. Espinoza, and B. Ren, “A high-resolution map of the three-dimensional chromatin interactome in human cells,” Nature 503, 290 (2013).

8 D. Leung, I. Jung, N. Rajagopal, A. Schmitt, S. Selvaraj, A. Y. Lee, C.-A. Yen, S. Lin, Y. Lin, Y. Qiu, et al., “Integrative analysis of haplotype-resolved epigenomes across human tissues,” Nature 518, 350 (2015).

9 N. H. Dryden, L. R. Broome, F. Dudbridge, N. Johnson, N. Orr, S. Schoenfelder, T. Nagano, S. Andrews, S. Wingett, I. Kozarewa, et al., “Unbiased analysis of potential targets of breast cancer susceptibility loci by capture Hi-C,” Genome Res. 24, gr–175034 (2014).

10 R. Jäger, G. Migliorini, M. Henrion, R. Kandaswamy, H. E. Speedy, A. Heindl, N. Whiffin, M. J. Carnicer, L. Broome, N. Dryden, et al., “Capture Hi-C identifies the chromatin interactome of colorectal cancer risk loci,” Nature Commun 6, 6178 (2015).

11 S. C. Baca, D. Prandi, M. S. Lawrence, J. M. Mosquera, A. Romanel, Y. Drier, K. Park, N. Kitabayashi, T. Y. MacDonald, M. Ghandi, et al., “Punctuated evolution of prostate cancer genomes,” Cell 153, 666–677 (2013).

12 M. Barbieri, M. Chotalia, J. Fraser, L.-M. Lavitas, J. Dostie, A. Pombo, and M. Nicodemi, “Complexity of chromatin folding is captured by the strings and binders switch model,” Proc. Natl. Acad. Sci. U. S. A. 109, 16173–16178 (2012).

13 A. L. Sanborn, S. S. P. Rao, S.-C. Huang, N. C. Durand, M. H. Huntley, A. I. Jewett, I. D. Bochkov, D. Chinnappan, A. Cutkosky, J. Li, K. P. Geeting, A. Gnirke, A. Melnikov, D. McKenna, E. K. Stamenova, E. S. Lander, and E. L. Aiden, “Chromatin extrusion explains key features of loop and domain formation in wild-type and engineered genomes,” Proc. Natl. Acad. Sci. USA 112, E6456–E6465 (2015).

14 G. Fudenberg, M. Imakaev, C. Lu, A. Goloborodko, N. Abdennur, and L. A. Mirny, “Formation of chromosomal domains by loop extrusion,” Cell Rep. 15, 2038–2049 (2016).

15 S. Bianco, D. G. Lupiáñez, A. M. Chiariello, C. Annunziatella, K. Kraft, R. Schöpflin, L. Wittler, G. Andrey, M. Vingron, A. Pombo, et al., “Polymer physics predicts the effects of structural variants on chromatin architecture,” Nat. Genetics 50, 662–667 (2018).

16 D. Jost, P. Carrivain, G. Cavalli, and C. Vaillant, “Modeling epigenome folding: formation and dynamics of topologically associated chromatin domains,” Nucleic Acids Res. 42, 9553–9561 (2014).

17 C. A. Brackley, J. M. Brown, D. Waithe, C. Babbs, J. Davies, J. R. Hughes, V. J. Buckle, and D. Marenduzzo, “Predicting the three-dimensional folding of cis-regulatory regions in mammalian genomes using bioinformatic data and polymer models,” Genome biology 17, 59 (2016).

18 S. Wang, J. Xu, and J. Zeng, “Inferential modeling of 3D chromatin structure,” Nucleic Acids Res. 43, e54 (2015).

19 P. Szalaj, P. J. Michalski, P. Wróblewski, Z. Tang, M. Kadlof, G. Mazzocco, Y. Ruan, and D. Plewczynski, “3D-GNOME: an integrated web service for structural modeling of the 3D genome,” Nucleic Acids Res. 44, W288 (2016).

20 H. Tjong, W. Li, R. Kalhor, C. Dai, S. Hao, K. Gong, Y. Zhou, H. Li, X. J. J. Zhou, M. A. Le Gros, C. A. Larabell, L. Chen, and F. Alber, “Population-based 3D genome structure analysis reveals driving forces in spatial genome organization.” Proc. Natl. Acad. Sci. USA 113, E1663–E1672 (2016).

21 M. Di Stefano, J. Paulsen, T. G. Lien, E. Hovig, and C. Micheletti, “Hi-C-constrained physical models of human chromosomes recover functionally-related properties of genome organization,” Sci. Rep. 6, 35985 (2016).

22 G. Shi, L. Liu, C. Hyeon, and D. Thirumalai, “Interphase Human Chromosome Exhibits Out of Equilibrium Glassy Dynamics,” Nat. Commun. 9, 3161 (2018).

23 S. Wang, J.-H. Su, B. J. Beliveau, B. Bintu, J. R. Moffitt, C.-t. Wu, and X. Zhuang, “Spatial organization of chromatin domains and compartments in single chromosomes,” Science 353, 598–602 (2016).

24 A. N. Boettiger, B. Bintu, J. R. Moffitt, S. Wang, B. J. Beliveau, G. Fudenberg, M. Imakaev, L. A. Mirny, C.-t. Wu, and X. Zhuang, “Super-resolution imaging reveals distinct chromatin folding for different epigenetic states,” Nature 529, 418–422 (2016).

25 C. Münkel and J. Langowski, “Chromosome structure predicted by a polymer model,” Phys. Rev. E 57, 5888–5896 (1998).

26 C. Münkel, R. Eils, S. Dietzel, D. Zink, C. Mehring, G. Wedemann, T. Cremer, and J. Langowski, “Compartmentalization of interphase chromosomes observed in simulation and experiment,” J. Mol. Biol. 285, 1053 – 1065 (1999).

27 M. Bohn, D. W. Heermann, and R. van Driel, “Random loop model for long polymers,” Phys. Rev. E. 76, 051805 (2007).

28 J. Mateos-Langerak, M. Bohn, W. de Leeuw, O. Giromus, E. M. M. Manders, P. J. Verschure, M. H. G. Indemans, H. J. Gierman, D. W. Heermann, R. van Driel, and S. Goetze, “Spatially confined folding of chromatin in the interphase nucleus,” Proc. Natl. Acad. Sci. USA 106, 3812–3817 (2009).

29 A. Hofmann and D. W. Heermann, “The role of loops on the order of eukaryotes and prokaryotes,” FEBS Letters 589, 2958–2965 (2015).

30 J. Fraser, C. Ferrai, A. M. Chiariello, M. Schueler, T. Rito, G. Laudanno, M. Barbieri, B. L. Moore, D. C. Kraemer, S. Aitken, et al., “Hierarchical folding and reorganization of chromosomes are linked to transcriptional changes in cellular differentiation,” Mol. Syst. Biol. 11, 852 (2015).

31 C. A. Brackley, J. Johnson, S. Kelly, P. R. Cook, and D. Marenduzzo, “Simulated binding of transcription factors to active and inactive regions folds human chromosomes into loops, rosettes and topological domains,” Nucleic Acids Res. 44, 3503–3512 (2016).

32 A. Buckle, C. A. Brackley, S. Boyle, D. Marenduzzo, and N. Gilbert, “Polymer simulations of heteromorphic chromatin predict the 3D folding of complex genomic loci,” Molecular Cell 72, 786 – 797.e11 (2018).

33 A. M. Chiariello, C. Annunziatella, S. Bianco, A. Esposito, and M. Nicodemi, “Polymer physics of chromosome large-scale 3D organisation,” Sci. Rep. 6, 29775 (2016).

34 M. Di Pierro, B. Zhang, E. L. Aiden, P. G. Wolynes, and J. N. Onuchic, “Transferable model for chromosome architecture,” Proc. Natl. Acad. Sci. USA 113, 12168–12173 (2016).

35 L. Liu, G. Shi, D. Thirumalai, and C. Hyeon, “Chain organization of human interphase chromosome determines the spatiotemporal dynamics of chromatin loci,” PLOS Comput. Biol. 14, 1–20 (2018).

36 L. Giorgetti, R. Galupa, E. Nora, T. Piolot, F. Lam, J. Dekker, G. Tiana, and E. Heard, “Predictive polymer modeling reveals coupled fluctuations in chromosome conformation and transcription,” Cell 157, 950 – 963 (2014).

37 G. Gürsoy, Y. Xu, A. L. Kenter, and J. Liang, “Computational construction of 3D chromatin ensembles and prediction of functional interactions of alpha-globin locus from 5C data,” Nucleic Acids Res. 45, 11547–11558 (2017).

38 G. Zhu, W. Deng, H. Hu, R. Ma, S. Zhang, J. Yang, J. Peng, T. Kaplan, and J. Zeng, “Reconstructing spatial organizations of chromosomes through manifold learning,” Nucleic Acids Res. 46, e50 (2018).

39 Q. Li, H. Tjong, X. Li, K. Gong, X. J. Zhou, I. Chiolo, and F. Alber, “The three-dimensional genome organization of *Drosophila melanogaster* through data integration,” Genome Biol. 18, 145 (2017).

40 F. Serra, M. Di Stefano, Y. G. Spill, Y. Cuartero, M. Goodstadt, D. Baù, and M. A. Marti-Renom, “Restraint-based threedimensional modeling of genomes and genomic domains,” FEBS Lett. 589, 2987–2995 (2015).

41 P. M. Goldbart and A. Zippelius, “Amorphous solid state of vulcanized macromolecules: a variational approach,” Phys. Rev. Lett. 71, 2256 (1993).

42 M. P. Solf and T. A. Vilgis, “Statistical mechanics of macro-molecular networks without replicas,” J. Phys. A: Mathematical and General 28, 6655 (1995).

43 J. D. Bryngelson and D. Thirumalai, “Internal constraints induce localization in an isolated polymer molecule,” Phys. Rev. Lett. 76, 542–545 (1996).

44 R. Zwanzig, “Effect of close contacts on the radius of gyration of a polymer,” J. Chem. Phys. 106, 2824–2827 (1997).

45 A. Cacciuto and E. Luijten, “Self-avoiding flexible polymers under spherical confinement,” Nano Lett. 6, 901–905 (2006).

46 H. Kang, Y.-G. Yoon, D. Thirumalai, and C. Hyeon, “Confinement-induced glassy dynamics in a model for chromosome organization,” Phys. Rev. Lett. 115, 198102 (2015).

47 J. R. Dixon, S. Selvaraj, F. Yue, A. Kim, Y. Li, Y. Shen, M. Hu, J. S. Liu, and B. Ren, “Topological domains in mammalian genomes identified by analysis of chromatin interactions,” Nature 485, 376–380 (2012).

48 B. Bonev, N. M. Cohen, Q. Szabo, L. Fritsch, G. L. Papadopoulos, Y. Lubling, X. Xu, X. Lv, J.-P. Hugnot, A. Tanay, and G. Cavalli, “Multiscale 3D genome rewiring during mouse neural development,” Cell 171, 557 – 572.e24 (2017).

49 Y. Zhang and D. W. Heermann, “Loops determine the mechanical properties of mitotic chromosomes,” PLoS One 6, 1–13 (2011).

50 G. Li, X. Ruan, R. Auerbach, K. Sandhu, M. Zheng, P. Wang, H. Poh, Y. Goh, J. Lim, J. Zhang, H. Sim, S. Peh, F. Mulawadi, C. Ong, Y. Orlov, S. Hong, Z. Zhang, S. Landt, D. Raha, G. Euskirchen, C.-L. Wei, W. Ge, H. Wang, C. Davis, K. I. Fisher-Aylor, A. Mortazavi, M. Gerstein, T. Gingeras, B. Wold, Y. Sun, M. Fullwood, E. Cheung, E. Liu, W.-K. Sung, M. Snyder, and Y. Ruan, “Extensive promoter-centered chromatin interactions provide a topological basis for transcription regulation,” Cell 148, 84–98 (2012).

51 Z. Tang, O. Luo, X. Li, M. Zheng, J. Zhu, P. Szalaj, P. Trzaskoma, A. Magalska, J. Wlodarczyk, B. Ruszczycki, P. Michalski, E. Piecuch, P. Wang, D. Wang, S. Tian, M. Penrad-Mobayed, L. Sachs, X. Ruan, C.-L. Wei, E. Liu, G. Wilczynski, D. Plewczynski, G. Li, and Y. Ruan, “CTCF-mediated human 3D genome architecture reveals chromatin topology for transcription,” Cell 163, 1611–1627 (2015).

52 D. Baù, A. Sanyal, B. R. Lajoie, E. Capriotti, M. Byron, J. B. Lawrence, J. Dekker, and M. A. Marti-Renom, “The threedimensional folding of the *α*-globin gene domain reveals formation of chromatin globules,” Nat. Struct. Mol. Biol. 18, 107–114 (2011).

53 C. Peng, L.-Y. Fu, P.-F. Dong, Z.-L. Deng, J.-X. Li, X.-T. Wang, and H.-Y. Zhang, “The sequencing bias relaxed characteristics of Hi-C derived data and implications for chromatin 3D modeling,” Nucleic Acids Res. 41, e183 (2013).

54 B. Friedman and B. O’Shaughnessy, “Short time behavior and universal relations in polymer cyclization,” J. Phys. II 1, 471–486 (1991).

55 C. Hyeon and D. Thirumalai, “Kinetics of interior loop formation in semiflexible chains,” J. Chem. Phys. 124, 104905 (2006).

56 N. Toan, P. Greg Morrison, C. Hyeon, and D. Thirumalai, “Kinetics of Loop Formation in Polymer Chains,” J. Phys. Chem. B 112, 6094–6106 (2008).

57 P. G. de Gennes, Scaling Concepts in Polymer Physics (Cornell University Press, Ithaca and London, 1979).

58 P. J. Flory, “The Configuration of Real Polymer Chains,” J. Chem. Phys. 17, 303–310 (1949).

59 A. Y. Grosberg and A. R. Khokhlov, Statistical Physics of Macromolecules (AIP Press, New York, 1994).

60 J. D. Halverson, J. Smrek, K. Kremer, and A. Y. Grosberg, “From a melt of rings to chromosome territories: the role of topological constraints in genome folding,” Rep. Prog. Phys. 77, 022601 (2014).

61 L. Liu and C. Hyeon, “Contact statistics highlight distinct organizing principles of proteins and rna,” Biophys. J. 110, 2320–2327 (2016).

62 A. Mortazavi, B. A. Williams, K. McCue, L. Schaeffer, and B. Wold, “Mapping and quantifying mammalian transcriptomes by RNA-Seq,” Nature Methods 5, 621–628 (2008).

63 J. Ernst, P. Kheradpour, T. S. Mikkelsen, N. Shoresh, L. D. Ward, C. B. Epstein, X. Zhang, L. Wang, R. Issner, M. Coyne, M. Ku, T. Durham, M. Kellis, and B. E. Bernstein, “Mapping and analysis of chromatin state dynamics in nine human cell types,” Nature 473, 43–49 (2011).

64 S. Goetze, J. Mateos-Langerak, H. J. Gierman, W. de Leeuw, O. Giromus, M. H. G. Indemans, J. Koster, V. Ondrej, R. Versteeg, and R. van Driel, “The three-dimensional structure of human interphase chromosomes is related to the transcriptome map,” Mol. Cell. Biol. 27, 4475–4487 (2007).

65 M. Levine, C. Cattoglio, and R. Tjian, “Looping back to leap forward: Transcription enters a new era,” Cell 157, 13 – 25 (2014).

66 C. Trapnell, B. A. Williams, G. Pertea, A. Mortazavi, G. Kwan, M. J. van Baren, S. L. Salzberg, B. J. Wold, and L. Pachter, “Transcript assembly and quantification by RNA-Seq reveals unannotated transcripts and isoform switching during cell differentiation,” Nature Biotechnol. 28, 511–515 (2010).

67 The ENCODE Project Consortium, “An Integrated Encyclopedia of DNA Elements in the Human Genome,” Nature 489, 57–74 (2012).

68 D. Vernimmen, F. Marques-Kranc, J. A. Sharpe, J. A. Sloane-Stanley, W. G. Wood, H. A. C. Wallace, A. J. H. Smith, and D. R. Higgs, “Chromosome looping at the human *α*-globin locus is mediated via the major upstream regulatory element (HS-40),” Blood 114, 4253–4260 (2009).

69 A. Astashyn, A. Shkeda, B. Rajput, C. M. Farrell, C. Wallin, D. Webb, D. R. Maglott, F. Thibaud-Nissen, G. R. Brown, H. Sun, J. M. Ostell, J. Weber, J. Hart, K. M. McGarvey, L. D. Riddick, M. J. Landrum, M. DiCuccio, M. R. Murphy, N. A. O’Leary, O. Ermolaeva, P. Tamez, P. Kitts, R. E. Tully, S. H. Rangwala, S. Pujar, S. M. Hiatt, T. D. Murphy, W. Wu, and K. D. Pruitt, “RefSeq: an update on mammalian reference sequences,” Nucleic Acids Res. 42, D756–D763 (2014).

70 A. Buckle, R.-s. Nozawa, D. A. Kleinjan, and N. Gilbert, “Functional characteristics of novel pancreatic Pax6 regulatory elements,” Hum. Mol. Genet. 27, 3434–3448 (2018).

71 A. Schedl, A. Seawright, D. A. Kleinjan, R. A. Quinlan, S. Danes, and V. van Heyningen, “Aniridia-associated translocations, DNase hypersensitivity, sequence comparison and transgenic analysis redefine the functional domain of PAX6,” Hum. Mol. Genet. 10, 2049–2059 (2001).

72 S. Bhatia, H. Bengani, M. Fish, A. Brown, M. Divizia, R. deÂa-Marco, G. Damante, R. Grainger, V. vanÂaHeyningen, and D. Kleinjan, “Disruption of Autoregulatory Feedback by a Mutation in a Remote, Ultraconserved PAX6 Enhancer Causes Aniridia,” Am. J. Hum. Genet. 93, 1126–1134 (2013).

73 M. Fritsche, S. Li, D. W. Heermann, and P. A. Wiggins, “A model for Escherichia coli chromosome packaging supports transcription factor-induced DNA domain formation,” Nucleic Acids Res. 40, 972–980 (2012).

74 M. Di Stefano, A. Rosa, V. Belcastro, D. di Bernardo, and C. Micheletti, “Colocalization of coregulated genes: A steered molecular dynamics study of human chromosome 19,” PLOS Comput. Biol. 9, 1–13 (2013).

75 D. Meluzzi and G. Arya, “Efficient estimation of contact probabilities from inter-bead distance distributions in simulated polymer chains,” J. Phys. Condens. Matter 27, 064120 (2015).

76 G. Fudenberg and M. Imakaev, “FISH-ing for captured contacts: towards reconciling FISH and 3C,” Nature Methods 14, 673–678 (2017).

77 Aronovitz, J.A. and Nelson, D.R., “Universal features of polymer shapes,” J. Phys. (Paris) 47, 1445–1456 (1986).

78 C. Hyeon, R. I. Dima, and D. Thirumalai, “Size, shape, and flexibility of rna structures,” J. Chem. Phys. 125, 194905 (2006).

79 C. H. Rycroft, “Voro++: A three-dimensional voronoi cell library in C++,” Chaos 19, 041111 (2009).

