## Supplementary Information for "Heterogeneous loop model to infer 3D chromosome structures from Hi-C"

### Hi-C data preparation

The contact frequency matrix obtained from a population of cells, the raw data of which are listed in Table 1, was first normalized by using the Knight-Ruiz (KR) method[1], so that the sum of each row and column of the matrix is unity. We then rescaled the KR-normalized matrix so that it satisfies $P(s) = 1$ at $s = 1$, and used it as the input contact probability matrix $\mathcal{P}$. Since genomic loci have different coordinates in sequence database for different genome assemblies, if necessary, we converted the genomic coordinates of human and mouse by choosing Hg19 and mm10, respectively, as their references.

### Pairwise contact probability in heterogeneous loop model

Similar to RLM[2], the harmonic-restraining energy potential of HLM for a chain of $N$ monomers can be written

$$U_{\mathcal{K}}(\mathbf{r}) = \frac{3}{2}\mathbf{r}^T\mathbf{K}\mathbf{r}, \tag{S1}$$

where $\mathbf{r} = (\vec{r}_1, \vec{r}_2, \cdots, \vec{r}_{N-1})^T$ and the translational degrees of freedom was removed by setting $\vec{r}_0 = (0, 0, 0)$. $\mathbf{K}$ is the Kirchhoff matrix:

$$\begin{pmatrix} \sum_{j=0,j\neq 1}^{N-1} k_{1j} & -k_{12} & \cdots & -k_{1,N-1} \\ -k_{21} & \sum_{j=0,j\neq 2}^{N-1} k_{2j} & \cdots & -k_{2,N-1} \\ \vdots & \vdots & \ddots & \vdots \\ -k_{N-1,1} & -k_{N-1,2} & \cdots & \sum_{j=0,j\neq N-1}^{N-1} k_{N-1,j} \end{pmatrix}. \tag{S2}$$

Then, the probability density of the distance between the $i$ and $j$-th monomer ($i < j$) projected on one dimension is

$$\begin{aligned} P(x_{ij}; \gamma_{ij}) &= \langle \delta[x_{ij} - (x_i - x_j)] \rangle \\ &= \int dx_1 \cdots dx_{N-1} \delta\left[x_{ij} - (x_i - x_j)\right] P(\mathbf{x}) \\ &\propto \int_0^\infty dq e^{iqx_{ij}} \int dx_1 \cdots dx_{N-1} e^{-iq(x_i - x_j)} e^{-\frac{1}{2}\mathbf{x}^T\mathbf{K}\mathbf{x}} \\ &\propto \int_0^\infty dq e^{iqx_{ij}} e^{-\frac{q^2}{4\gamma_{ij}}} \\ &\propto e^{-\gamma_{ij} x_{ij}^2}, \end{aligned} \tag{S3}$$

where we have used $\langle \exp\left(\sum_n \xi_n x_n\right) \rangle = \exp\left(\frac{1}{2}\sum_{nm}(K^{-1})_{nm}\xi_n\xi_m\right)$. The value of $\gamma_{ij}$ depends on the topology of 'vulcanized' polymer chain, dictated by $\mathbf{K}$ matrix, and is related with the covariance between the positions of $i$ and $j$-th monomer $\sigma_{ij} = \langle \delta\vec{r}_i \cdot \delta\vec{r}_j \rangle$ as follows

$$\gamma_{ij} = \begin{cases} \frac{1}{2(\sigma_{ii} + \sigma_{jj} - 2\sigma_{ij})}, & i > 0 \\ \frac{1}{2\sigma_{jj}}, & i = 0 \end{cases} \tag{S4}$$

where $\sigma_{ij} (= (\Sigma)_{ij})$ is the elements of inverse matrix $\Sigma = \mathbf{K}^{-1}$. Finally, the probability density of pairwise distance in 3D is[2]

$$P(r_{ij}; \gamma_{ij}) = 4\gamma_{ij}^{3/2}/\sqrt{\pi} r_{ij}^2 e^{-\gamma_{ij} r_{ij}^2}. \tag{S5}$$

Typical profiles of $P(r_{ij}; \gamma_{ij})$ for varying $\gamma_{ij}$ are shown in Fig. S1A with the relation between $p_{ij}$ and $\gamma_{ij}$ given in the inset.

Eq. S5 enables us to evaluate a few quantities of interest directly. For a pair of monomers in contact with the condition of $r_{ij} < r_c$, the pairwise contact probability, $p_{ij}$, is given by

$$\begin{aligned} p_{ij} &= \int_0^{r_c} P(r_{ij}) dr_{ij} \\ &= \mathrm{erf}(\gamma_{ij}^{1/2} r_c) - 2r_c\sqrt{\frac{\gamma_{ij}}{\pi}} e^{-\gamma_{ij} r_c^2}, \end{aligned} \tag{S6}$$

with $\mathrm{erf}(x) = \frac{2}{\sqrt{\pi}}\int_0^x dt e^{-t^2}$. In addition, the mean pairwise distance is

$$\langle r_{ij} \rangle = \int_0^\infty r_{ij} P(r_{ij}) dr_{ij} = \frac{2}{\sqrt{\pi}\gamma_{ij}^{1/2}}, \tag{S7}$$

and the mean square distance equals to

$$\langle r_{ij}^2 \rangle = \int_0^\infty r_{ij}^2 P(r_{ij}) dr_{ij} = \frac{3}{2\gamma_{ij}}. \tag{S8}$$

For $\gamma_{ij} r_c^2 (\equiv g_{ij}) \ll 1$ (or $\langle r_{ij} \rangle \gg r_c$), $p_{ij}$ is approximated as

$$\begin{aligned} p_{ij} &= \frac{2}{\sqrt{\pi}}\left\{ \int_0^{g_{ij}^{1/2}} [1 - t^2 + \mathcal{O}(t^4)] dt - g_{ij}^{1/2}[1 - g_{ij} + \mathcal{O}(g_{ij}^2)] \right\} \\ &= \frac{1}{3\sqrt{\pi}} g_{ij}^{3/2} + \mathcal{O}(g_{ij}^{5/2}). \end{aligned} \tag{S9}$$

Replacing $g_{ij}$ with $\langle r_{ij} \rangle$ using Eq. S7, one obtains a scaling relation between $p_{ij}$ and $\langle r_{ij} \rangle$ as

$$p_{ij} \sim \left[\frac{4r_c^2}{\pi\langle r_{ij}\rangle^2}\right]^{3/2} \sim \langle r_{ij} \rangle^{-3}. \tag{S10}$$

As shown in Fig. S1B, the scaling $p_{ij}^{-1} \sim \langle r_{ij} \rangle^3$ holds for large $\langle r_{ij} \rangle$.

RLM was developed to understand the scaling of the spatial distance between two genomic loci with respect to their genomic distance[2]. The original RLM assumes that all loops have the same interaction strength (i.e., $k_{ij} = 3$ or $0$ $k_BT/a^2$), and the overall compactness of the chain was adjusted by the total number of loops ($N_l = \sum_{i>j} \delta(k_{ij} - 3)$). Therefore, any quantity of interest, e.g., $P(r_{ij})$, needs to be averaged over different instances of $\mathcal{K}$ with the same value of $N_l$. In the worst case, this requires $2^{(N-1)(N-2)/2}$ implementations of $\mathcal{K}$, which renders a precise evaluation of $P(r_{ij})$ impractical. In this study, instead of varying $N_l$, we relax the constraint on $k_{ij}$ so that it can take any non-negative value.

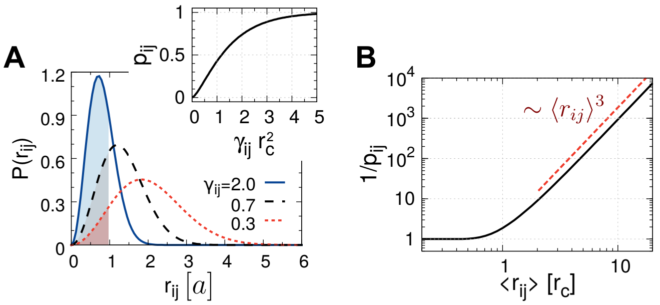


FIG. S1. (A) Probability density of pairwise distance $P(r_{ij})$ at various values of $\gamma_{ij}$ (Eq. S5), where the inset shows contact probability $p_{ij}$ as a function of $\gamma_{ij} r_c^2$ (Eq. S6). (B) Inverse of $p_{ij}$ is plotted as a function of the mean pairwise distance $\langle r_{ij}\rangle$ in a log-log scale, where it shows $p_{ij}^{-1} \sim \langle r_{ij}\rangle^3$ at $\langle r_{ij}\rangle \gg r_c$ (Eq. S10).

### Inferring interaction strengths by direct inversion

In HLM, $p_{ij}$ increases *monotonically* with $\gamma_{ij}$ at given $r_c$ (Fig. S1A), which allows one to determine the value of $\gamma_{ij}$ from $p_{ij}$. Given a contact probability matrix $\mathcal{P}$ of elements $p_{ij}$, we can further derive the interaction strength matrix $\mathcal{K}$ of elements $k_{ij}$ in three steps

$$\mathcal{P} \to \Sigma \to \mathbf{K} \to \mathcal{K}. \quad \text{(S11)}$$

More specifically, the interaction strength matrix $\mathcal{K}$ for HLM can be obtained from $\mathcal{P}$ via the following steps, which we call the *direct inversion*:

1. Construct the matrix $\Sigma$ using the relations of diagonal elements $\sigma_{ii} = \frac{1}{2\gamma_{0i}}$ for $i > 0$; and the off-diagonal elements $\sigma_{ij} = \frac{1}{2}\left(\sigma_{ii} + \sigma_{jj} - \frac{1}{2\gamma_{ij}}\right)$ for $i, j > 0, i \neq j$.
2. Invert the matrix ($\Sigma \to \Sigma^{-1}$) to get the Kirchhoff matrix $\mathbf{K}$.
3. Determine the interaction strengths $k_{ij} (= (\mathcal{K})_{ij})$ from (i) $k_{ij} = -(\mathbf{K})_{ij}$ for $i, j > 0, i \neq j$; (ii) $k_{0i} = \sum_j (\mathbf{K})_{ij}$ for $i, j > 0$.

However, even a small sampling error, if any, in the contact probability matrix $\mathcal{P}$ may result in unphysical values of $\mathcal{K}$ with this protocol. We demonstrate this issue clearly using molecular dynamics (MD) simulations of three toy models ($N = 20$) characterized with different intra-chain loops (i.e., different $\mathcal{K}$-matrix): (i) a chain with a single loop (Fig. S9); (ii) a chain with two nested loops (Fig. S10); (iii) a chain composed of two blocks of monomers without any inter-block attraction (Fig. S11). For the case of the single-loop polymer (model (i)), $\mathcal{P}$ estimated based on the conformational ensemble from MD simulation resembles $\mathcal{P}^*$, where the superscript $^*$ denotes the *true* value, with a small relative error of 0.018; however, $\mathcal{K}$ obtained from $\mathcal{P}$ using the aforementioned direct inversion gives rise to unphysical matrix elements $k_{ij} < 0$. The same issue ($k_{ij} < 0$) was encountered for the two other cases (models (ii) and (iii)). For the three toy models, we circumvented the issue of $k_{ij} < 0$ resulting from the direct inversion through the constrained optimization, which is explained in the METHOD section in the main text and illustrated along the magenta arrows in Figs. S9-S11.

We note that Hi-C data is still an outcome of sampling over finite number of cell population, and hence a small but finite amount of error is inevitably included in Hi-C data. The same issue arises when the direct inversion is applied to Hi-C. Therefore, in the framework of HLM, we use the constrained optimization to determine $\widetilde{\mathcal{K}}$-matrix that can serve as a proxy of $\mathcal{K}$, which are followed by MD simulations using HLM potential.

### Molecular dynamics simulations

To produce a conformational ensemble of chromosome using HLM via the enhanced sampling, we performed the low-friction Langevin simulations by numerically integrating the following equation[3],

$$m\frac{d^2\vec{r}_i}{dt^2} = -\zeta_{\text{MD}}\frac{d\vec{r}_i}{dt} - \vec{\nabla}_{\vec{r}_i} U(\vec{r}_1, \vec{r}_2, \ldots) + \vec{\xi}(t), \quad \text{(S12)}$$

We chose a friction coefficient $\zeta_{\text{MD}} = 1.0m/\tau_{\text{MD}}$ and a time step $\delta t = 0.01\tau_{\text{MD}}$ with the characteristic time scale $\tau_{\text{MD}} = (ma^2/\epsilon)^{1/2}$. The whole simulation was carried out with three steps. (i) Random Gaussian chains were first equilibrated for 500 $\tau_{\text{MD}}$ under the energy potential $U_{\mathcal{K}}(\mathbf{r})$ without nonbonded interaction term. At this stage, excluded volume interaction is absent. (ii) The simulations were performed under the full HLM potential $U_{\text{HLM}}(\mathbf{r})$ for 100 $\tau_{\text{MD}}$ but with extra care. Excessive overlaps between monomers generated from the foregoing stage, were eliminated by gradually increasing the contribution from the short-range repulsive potential, which was achieved by using the LJ potential term $u_{\text{LJ}}(r_{ij}) = \min\{u_c, u_{\text{LJ}}(r_{ij})\}$ with gradually increasing $u_c$. (iii) The production runs were generated for $5 \times 10^5\tau_{\text{MD}}$, during which chain configurations were collected every 50 $\tau_{\text{MD}}$. Simulations were all carried out by using ESPResSo 3.3.1 package[4]. For any chromosome ensemble discussed in this work, it took less than 5 hours on a single CPU to generate them.

### Quality of HLM-generated structures

Since the only input of HLM is the contact probability matrix from Hi-C, the quality of resulting structures depends on the accuracy of input matrix $\mathcal{P}$. For given Hi-C data acquired from an experiment with a finite number of samples, if we were to increase the resolution of the model by reducing the genomic size of each monomer of HLM, the number of contacts counted for each pair of monomers decreases, which results in a less precise $\mathcal{P}$. Therefore, chromosome model generated from HLM or consequently $\widetilde{\mathcal{P}}$ becomes less accurate for a noisy input $\mathcal{P}$. As listed in Table 1, the Pearson correlation between HLM and Hi-C decreases as we either try to increase the resolution of model or decrease the number of sampled cells. Furthermore, to make tractable the parametrization of $\widetilde{\mathcal{K}}$ matrix via the algorithm of constrained optimization (Eq. 9), $N$ should not be too large.

### Clustering analysis on chromosomes with conformational variability

To characterize the variability in the conformational ensemble in Fig. 6D quantitatively, we cluster the HLM-generated structures using hierarchical clustering algorithm. We first defined three domains labeled as "L", "M" and "N" based on the sign of the first principle component of $\mathcal{Z}$ matrix (see the left tracks of Fig. 6B). Structures were hierarchically clustered according to structural similarity assessed by the distance-based root-mean-square deviation (dRMSD). For any two structures, say $\alpha$ and $\beta$, their similarity was measured by

$$\mathrm{dRMSD}_{\alpha,\beta} = \sqrt{\sum_{\{\text{X-Y}\}} \frac{1}{3}(r^{\alpha}_{\text{X-Y}} - r^{\beta}_{\text{X-Y}})^2}, \qquad \text{(S13)}$$

where $r_{\text{X-Y}}$ is the distance between the geometric centers of two different domains X and Y (X,Y $\in$ {L,M,N}). The dendrogram depicted in Fig. S8A identifies *at least* 4 main classes of conformations. Chromatins fold into compact globules in the class-1, but adopt elongated conformations in the class-4. The class-2 and -3 can be identified separately from the class-1 and -4 in the 2D phase plane drawn as a function of $r_{\text{L-M}}$ and $r_{\text{L-N}}$ (Fig. S8B). Since the structural interconversion among different chromosome conformations is an unusually time-consuming, glass-like process[5], the contact probability matrix $\widetilde{\mathcal{P}}$ (Fig. 6E) is in effect an outcome of *quenched*-average[2] over distinct conformations.

### Quantifying the similarity between contact probabilities from Hi-C and modeling

It has been recently proposed[6] that compared with a global Pearson or Spearman correlation coefficient, the reproducibility of Hi-C data can be better assessed by measuring the Pearson correlation PC($s$), at each value of genomic separation $s$, which minimizes the dependence of contact frequency on the genomic distance (e.g., see Fig. S7E). A underlying assumption for this quantity is that the mean contact probability as a function of genomic distance, $P(s)$, does not change too much. Whereas this is probably true for different replicates in Hi-C experiment, it is not guaranteed for modeling.

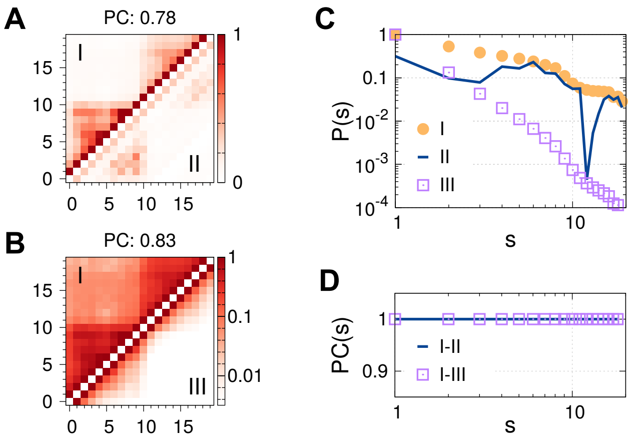


FIG. S2. Similarity between contact probablity matrices. Comparison of contact probability matrices I and II (A), I and III (B), with the overall Pearson correlations labeled on top. (C) Mean contact probability, (D) Pearson correlation as a function of genomic distance.

Two examples are shown in Fig. S2A and B. Although the contact probability matrix I and II (III) are perfectly correlated at each genomic scale (Fig. S2D), their overall similarity is low due to the different profiles of $P(s)$ (Fig. S2C). Therefore, even though PC($s$) is indeed more informative than an overall correlation coefficient, we chose the latter to quantify the similarity of contact probabilities between Hi-C and our model.

[1] P. A. Knight and D. Ruiz, "A fast algorithm for matrix balancing," IMA J. Numer. Anal. **33**, 1029 (2013).
[2] M. Bohn, D. W. Heermann, and R. van Driel, "Random loop model for long polymers," Phys. Rev. E. **76**, 051805 (2007).
[3] T. Veitshans, D. Klimov, and D. Thirumalai, "Protein folding kinetics: timescales, pathways and energy landscapes in terms of sequence-dependent properties," Folding Des. **2**, 1–22 (1997).
[4] H. J. Limbach, A. Arnold, B. A. Mann, and C. Holm, "ESPResSo – an extensible simulation package for research on soft matter systems," Comput. Phys. Commun. **174**, 704–727 (2006).
[5] G. Shi, L. Liu, C. Hyeon, and D. Thirumalai, "Interphase Human Chromosome Exhibits Out of Equilibrium Glassy Dynamics," Nat. Commun. **9**, 3161 (2018).
[6] T. Yang, F. Zhang, G. G. Yardimci, F. Song, R. C. Hardison, W. S. Noble, F. Yue, and Q. Li, "HiCRep: assessing the reproducibility of Hi-C data using a stratum-adjusted correlation coefficient," Genome Res. **27**, 1939–1949 (2017).
[7] J. Mateos-Langerak, M. Bohn, W. de Leeuw, O. Giromus, E. M. M. Manders, P. J. Verschure, M. H. G. Indemans, H. J. Gierman, D. W. Heermann, R. van Driel, and S. Goetze, "Spatially confined folding of chromatin in the interphase nucleus," Proc. Natl. Acad. Sci. USA **106**, 3812–3817 (2009).
[8] S. Wang, J.-H. Su, B. J. Beliveau, B. Bintu, J. R. Moffitt, C.-t. Wu, and X. Zhuang, "Spatial organization of chromatin domains and compartments in single chromosomes," Science **353**, 598–602 (2016).
[9] R. M. Kuhn, D. Haussler, and W. J. Kent, "The UCSC genome browser and associated tools," Brief Bioinform **14**, 144–161 (2013).
[10] G. Li, X. Ruan, R. Auerbach, K. Sandhu, M. Zheng, P. Wang, H. Poh, Y. Goh, J. Lim, J. Zhang, H. Sim, S. Peh, F. Mulawadi, C. Ong, Y. Orlov, S. Hong, Z. Zhang, S. Landt, D. Raha, G. Euskirchen, C.-L. Wei, W. Ge, H. Wang, C. Davis, K. I. Fisher-Aylor, A. Mortazavi, M. Gerstein, T. Gingeras, B. Wold,

Y. Sun, M. Fullwood, E. Cheung, E. Liu, W.-K. Sung, M. Snyder, and Y. Ruan, "Extensive promoter-centered chromatin interactions provide a topological basis for transcription regulation," Cell **148**, 84–98 (2012).

[11]Z. Tang, O. Luo, X. Li, M. Zheng, J. Zhu, P. Szalaj, P. Trzaskoma, A. Magalska, J. Wlodarczyk, B. Ruszczycki, P. Michalski, E. Piecuch, P. Wang, D. Wang, S. Tian, M. Penrad-Mobayed, L. Sachs, X. Ruan, C.-L. Wei, E. Liu, G. Wilczynski, D. Plewczynski, G. Li, and Y. Ruan, "CTCF-mediated human 3D genome architecture reveals chromatin topology for transcription," Cell **163**, 1611–1627 (2015).

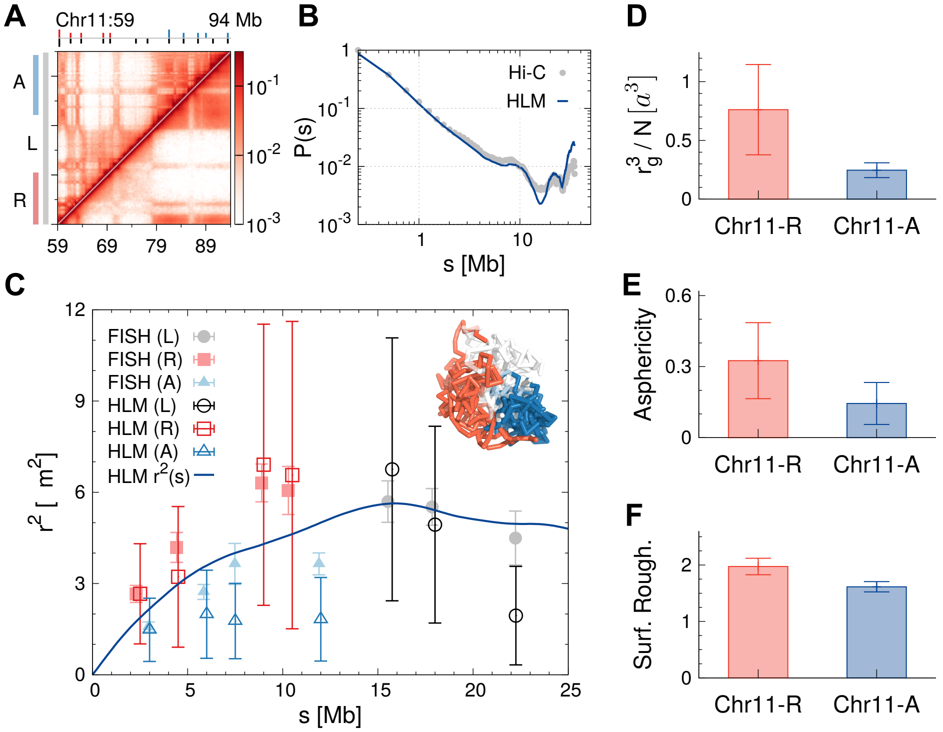


FIG. S3. A 35 Mb genomic region on chr11 in IMR90 cells modeled by HLM. Mateos-Langerak *et al.* have performed FISH experiment[7] in this region, with FISH probes distributed within a transcriptionally active ridge domain, inactive anti-ridge domain, and a longer region including both. They are labeled as "R","A" and "L", respectively. The genomic positions of the probes are labeled by sticks at the top of (A), below which is the heatmap of contact probabilities from Hi-C (upper diagonal region) and HLM (lower diagonal region). (B) Mean contact probability $P(s)$. (C) Pairwise square distance $r_{ij}^2$ between the FISH probes as a function of the genomic distance $s$, as well as the mean square distance $r^2(s) = \sum_{i=0}^{N-s-1} r_{i,i+s}^2/(N-s)$ (the solid line). Structural ensemble is illustrated with the ridge and anti-ridge domains colored red and blue, respectively. (D) Compactness, (E) asphericity, and (F) roughness of the surface of the domains.

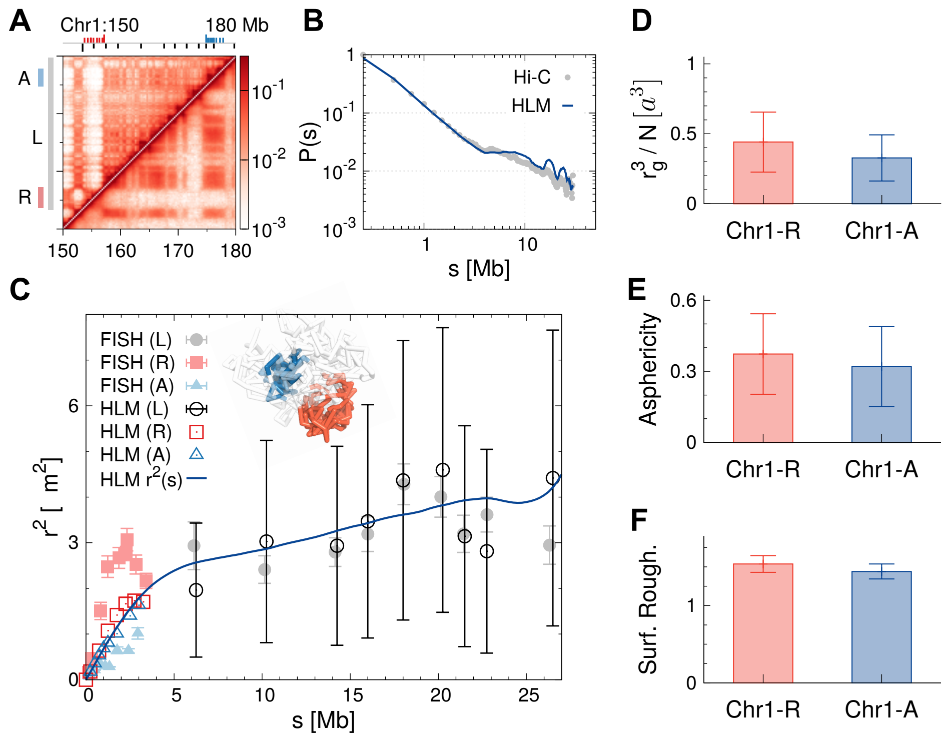


FIG. S4. A 30 Mb genomic region on chr1 in IMR90 cells modeled by HLM (see also the caption of Fig. S3).

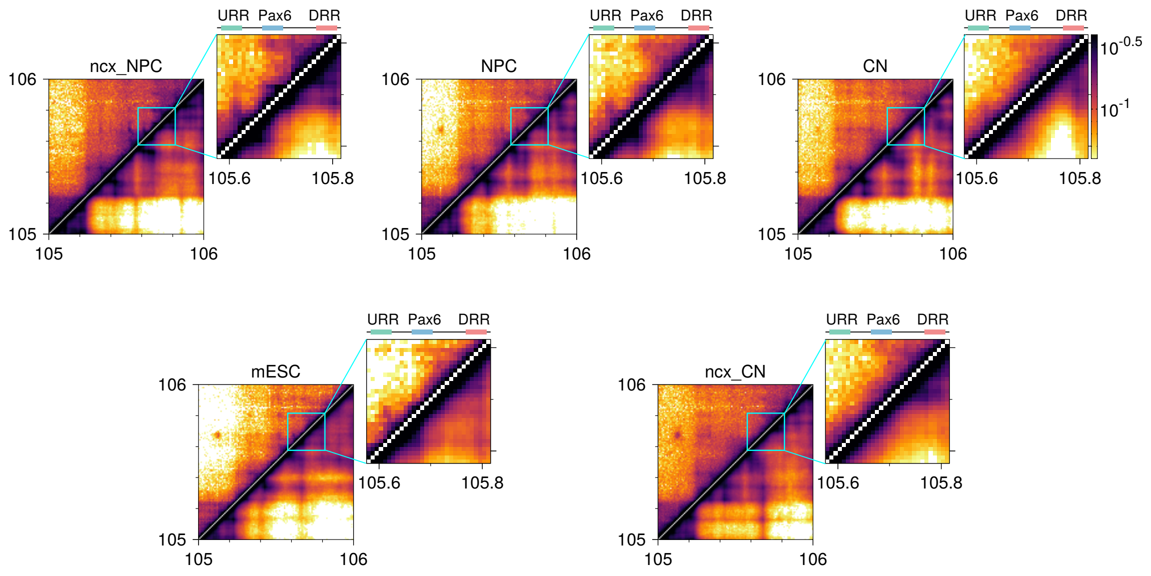


FIG. S5. Contact probabilities calculated from Hi-C (upper diagonal region) and HLM (lower diagonal region) around the mouse Pax6 gene for five different cell types. A 200 kb region was zoomed in, which highlights the contacts between the three "simulated" FISH probes (URR, Pax6, and DRR).

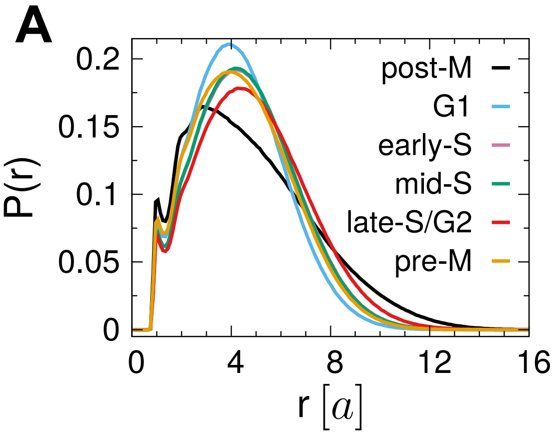


FIG. S6. (A) Probability densities of pairwise distance $P(r) = \frac{2}{N(N-1)} \sum_{i>j} \delta(r_{ij} - r)$ in chr19 of mESC at different phases of the cell cycle. (B) A zoom-in view of $P(r)$ at small values of $r$. Because of the elongated shape, the post-M phase has $P(r)$ with a fatter tail than other phases. In terms of local compactness, $P(r)$ at $r < 2a$ (Fig. S6B) suggests that post-M phase is the most compact, followed by pre-M phase and so forth (post-M > pre-M > G1 > early-S ≥ mid-S > late-S/G2), the order of which is identical to that assessed by the monomer volume ($\overline{v}$) (Fig. 5B).

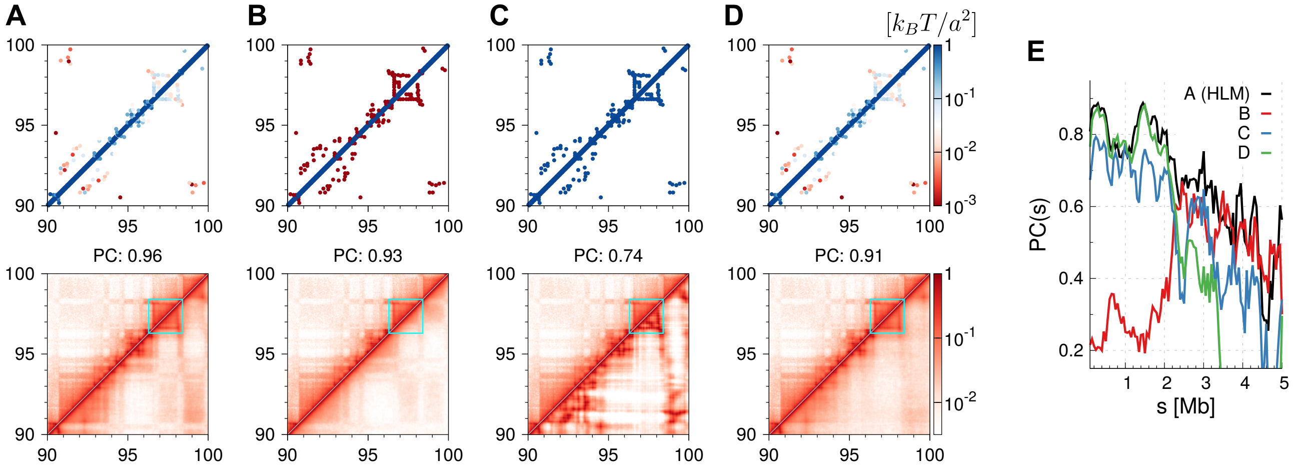


FIG. S7. Comparison of HLM with its variants by modeling a 10 Mb region of GM12878 cells. Four models are considered: multi-block copolymer (A,B,C) and homopolymer (D) model with harmonic springs of various (A,D) or uniform strengths (B,C). For each model, the top panel shows the corresponding spring strength matrix, and the resulting contact probabilities (lower diagonal) are contrasted with those measured by Hi-C (upper diagonal) at the bottom panel. A domain with edges showing enriched contacts is highlighted by a cyan box. (E) Pearson correlation of contact probabilities between Hi-C and modeling as a function of the genomic separation.

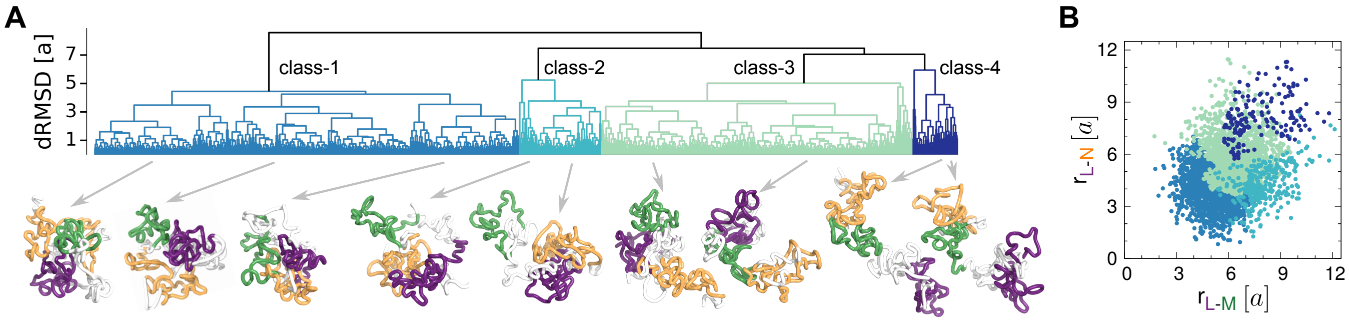


FIG. S8. Variability in the HLM-generated conformational ensemble for a 10 Mb-genomic region of chr5 in GM12878 cells (Fig. 6D). (A) Dendrogram of chromosome conformations from hierarchical clustering. Illustrated are the chromosome conformations from the four classes with L, M, N domains colored in purple, green, orange, respectively, following the domain labels assigned in Fig. 6B. (B) Scatter plot of inter-domain distances $r_{\text{L-M}}$ versus $r_{\text{L-N}}$ of structures in different classes.

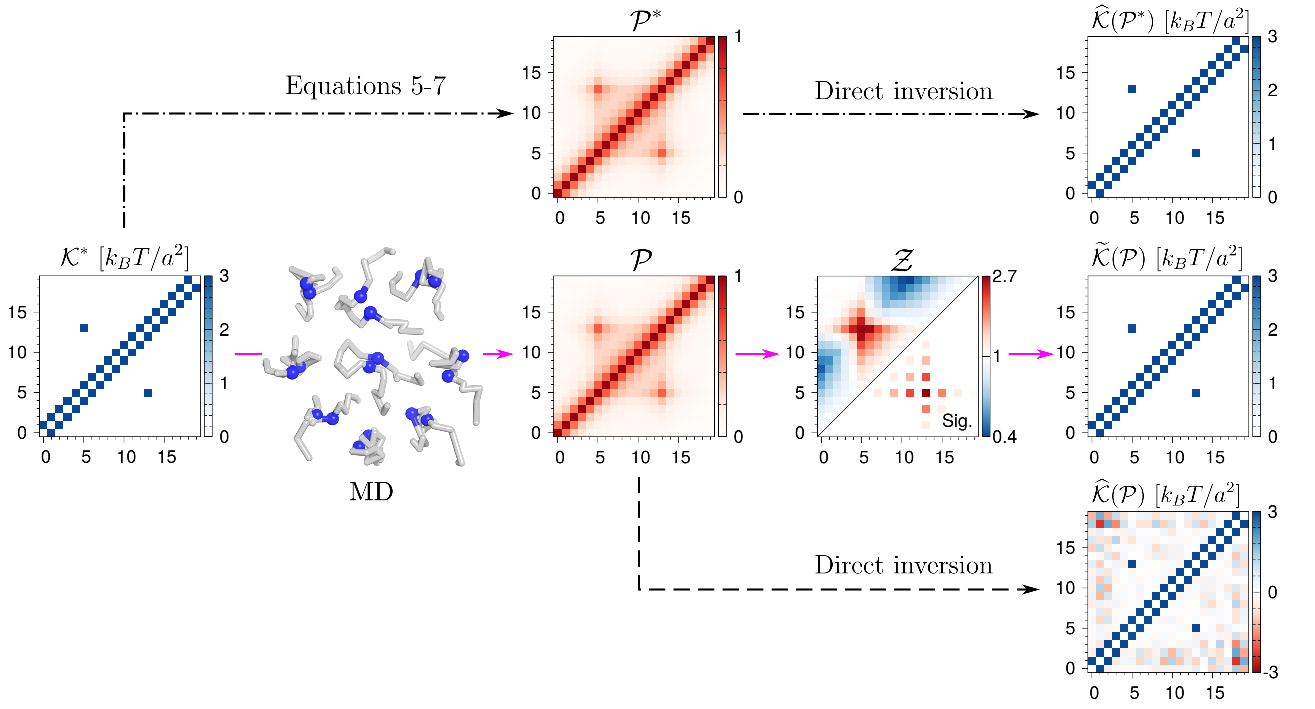


FIG. S9. HLM illustrated by a model of single-loop polymer chain. Starting from a true spring strength matrix $\mathcal{K}^*$, one can obtain either the contact probability matrix $\mathcal{P}^*$ analytically, or $\mathcal{P}$ (as an estimate of $\mathcal{P}^*$) numerically. Through the direct inversion (Eq. S11), $\mathcal{P}^*$ reproduces $\widehat{\mathcal{K}}(\mathcal{P}^*) = \mathcal{K}^*$, but $\mathcal{P}$, which is similar to $\mathcal{P}^*$ except for small deviation, generates $\widehat{\mathcal{K}}(\mathcal{P})$ that contains unphysical negative elements due to numeric errors. By contrast, $\mathcal{K}^*$ can be still inferred from $\mathcal{P}$, $\widetilde{\mathcal{K}}(\mathcal{P}) \approx \mathcal{K}^*$ with a relative error of $8 \times 10^{-4}$, using the constrained optimization (Eq. 9) on the significant contacts selected from $\mathcal{Z}$ (lower diagonal region).

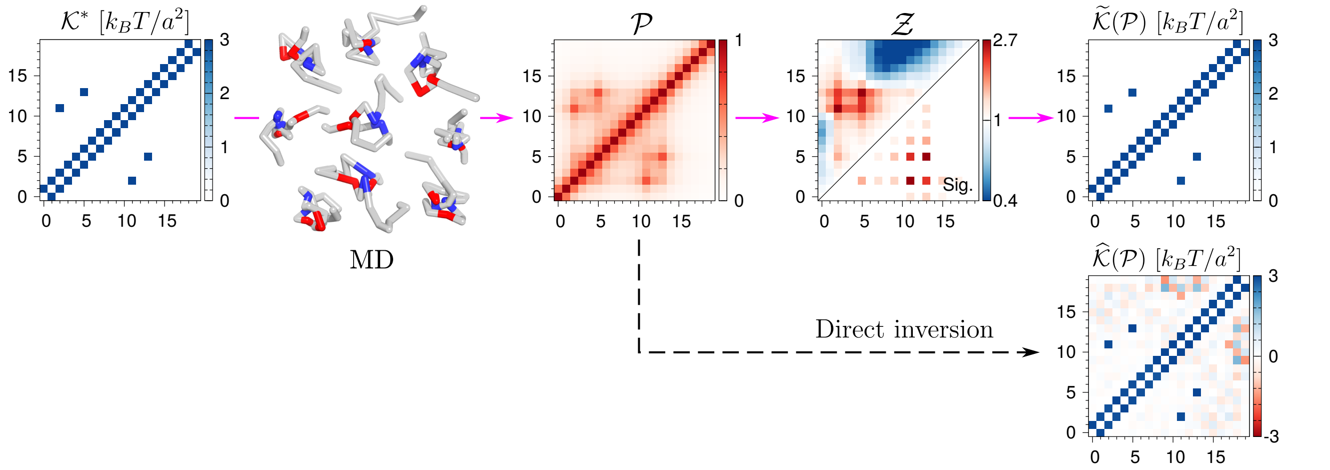


FIG. S10. HLM illustrated by a model of two-nested-loop polymer chain. The chain is composed of 20 monomers. One loops is anchored between the second and 11-th monomers, and the other is between the 5-th and 13-th monomers. In contrast to the direct inversion which generates unphysical negative elements in $\widehat{\mathcal{K}}(\mathcal{P})$, constrained optimization leads to $\widetilde{\mathcal{K}}(\mathcal{P})$, that is similar to $\mathcal{K}^*$ with a relative error of 0.002.

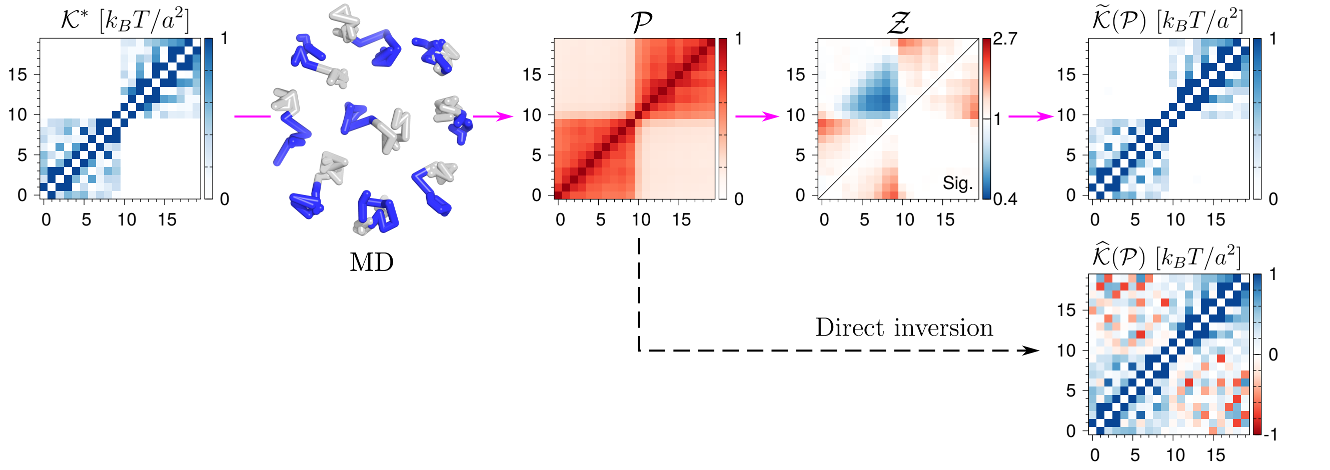


FIG. S11. HLM illustrated by a model of diblock copolymer chain. The chain is composed of 20 monomers. $\mathcal{K}^*$-matrix indicates the presence of heterogeneous intra-block attractions within the first $0 \leq i,j < 10$ and the second block $10 \leq i,j < 20$, but there is no such interaction between the two blocks. Compared to $\widetilde{\mathcal{K}}(\mathcal{P})$ calculated from direct inversion of $\mathcal{P}$-matrix which is fraught with many negative the inter-block interaction strength demonstrated in the off-diagonal block, $\widetilde{\mathcal{K}}(\mathcal{P})$ inferred by the constrained optimization display a better resemblance to $\mathcal{K}^*$ with a relative error of 0.044.

TABLE S1. TADs on chr21 of human IMR90 cells whose pairwise distances were measured in Ref.[8], and computed in Fig. 1.

| TAD index | Start(bp) | End(bp) | Center(bp) | $i$ | A/B |
|---|---|---|---|---|---|
| 2 | 13,280,000 | 16,160,000 | 14,720,000 | 2 | B |
| 3 | 16,160,000 | 18,280,000 | 17,220,000 | 12 | B |
| 4 | 18,320,000 | 21,080,000 | 19,700,000 | 22 | B |
| 5 | 21,240,000 | 23,160,000 | 22,200,000 | 32 | B |
| 6 | 24,320,000 | 25,680,000 | 25,000,000 | 44 | B |
| 7 | 26,160,000 | 27,000,000 | 26,580,000 | 50 | B |
| 8 | 27,040,000 | 28,000,000 | 27,520,000 | 54 | A |
| 9 | 28,000,000 | 29,320,000 | 28,660,000 | 58 | B |
| 10 | 29,360,000 | 29,680,000 | 29,520,000 | 62 | B |
| 11 | 29,760,000 | 31,360,000 | 30,560,000 | 66 | B |
| 12 | 31,400,000 | 31,960,000 | 31,680,000 | 70 | B |
| 13 | 31,960,000 | 32,600,000 | 32,280,000 | 73 | B |
| 14 | 32,600,000 | 32,920,000 | 32,760,000 | 75 | B |
| 15 | 33,080,000 | 33,800,000 | 33,440,000 | 77 | B |
| 16 | 33,840,000 | 34,200,000 | 34,020,000 | 80 | A |
| 17 | 34,200,000 | 34,680,000 | 34,440,000 | 81 | A |
| 18 | 34,680,000 | 34,920,000 | 34,800,000 | 83 | A |
| 19 | 34,920,000 | 36,440,000 | 35,680,000 | 86 | A |
| 20 | 36,440,000 | 36,640,000 | 36,540,000 | 90 | A |
| 21 | 36,680,000 | 37,440,000 | 37,060,000 | 92 | A |
| 22 | 37,440,000 | 37,880,000 | 37,660,000 | 94 | B |
| 23 | 37,960,000 | 38,720,000 | 38,340,000 | 97 | B |
| 24 | 38,720,000 | 39,760,000 | 39,240,000 | 100 | B |
| 25 | 39,760,000 | 41,440,000 | 40,600,000 | 106 | B |
| 26 | 41,440,000 | 42,120,000 | 41,780,000 | 111 | B |
| 27 | 42,120,000 | 42,840,000 | 42,480,000 | 113 | B |
| 28 | 42,840,000 | 43,120,000 | 42,980,000 | 115 | B |
| 29 | 43,160,000 | 44,040,000 | 43,600,000 | 118 | A |
| 30 | 44,040,000 | 44,360,000 | 44,200,000 | 120 | A |
| 31 | 44,360,000 | 45,040,000 | 44,700,000 | 122 | A |
| 32 | 45,040,000 | 45,400,000 | 45,220,000 | 124 | A |
| 33 | 45,440,000 | 46,160,000 | 45,800,000 | 127 | A |
| 34 | 46,160,000 | 46,944,323 | 46,552,161 | 130 | A |

$i$ is the index of the corresponding monomer in HLM. A/B type is determined by a principal component analysis of the inter-TAD distance matrix measured in the experiment.

TABLE S2. FISH probes in chr11 of human IMR90 cells whose pairwise distance were measured in Ref.[7], and computed in Fig. S3.

| Domain | $q$ | Start(bp) | End(bp) | Center(bp) | $i$ |
|---|---|---|---|---|---|
| R | 1 | 59,145,349 | 59,328,495 | 59,236,922 | 0 |
| | 0 | 61,446,617 | 61,635,131 | 61,540,874 | 10 |
| | 0 | 63,606,476 | 63,728,827 | 63,667,651 | 18 |
| | 0 | 68,035,816 | 68,234,792 | 68,135,304 | 36 |
| | 0 | 69,453,280 | 69,614,785 | 69,534,032 | 42 |
| A | 1 | 81,410,783 | 81,515,783 | 81,463,283 | 89 |
| | 0 | 84,330,302 | 84,491,603 | 84,410,952 | 101 |
| | 0 | 87,242,814 | 87,391,556 | 87,317,185 | 113 |
| | 0 | 88,840,806 | 89,052,495 | 88,946,650 | 119 |
| | 0 | 93,270,300 | 93,463,527 | 93,366,913 | 137 |
| L | 1 | 59,145,349 | 59,328,495 | 59,236,922 | 0 |
| | 0 | 74,678,334 | 74,845,650 | 74,761,992 | 63 |
| | 0 | 77,016,132 | 77,155,090 | 77,085,611 | 72 |
| | 0 | 81,410,783 | 81,515,783 | 81,463,283 | 89 |
| | 0 | 84,330,302 | 84,491,603 | 84,410,952 | 101 |
| | 0 | 87,242,814 | 87,391,556 | 87,317,185 | 113 |
| | 0 | 90,287,090 | 90,448,063 | 90,367,576 | 125 |
| | 0 | 93,270,300 | 93,463,527 | 93,366,913 | 137 |

In the experiment[7], distances were measured between the reference probe ($q = 1$) and other probes ($q = 0$) within a transcriptionally active ridge domain (R), a transcriptionally inactive anti-ridge domain (A), and a longer genomic region including both (L). The genomic positions of probes were lifted from hg15 to hg19[9]. $i$ is the index of the corresponding monomer in HLM.

TABLE S3. FISH probes in chr1 of human IMR cells. The pairwise distance were measured in Ref.[7], and computed in Fig. S4.

| Domain | $q$ | Start(bp) | End(bp) | Center(bp) | $i$ |
|---|---|---|---|---|---|
| R | 0 | 153,688,049 | 153,838,214 | 153,763,131 | 15 |
| | 0 | 154,258,113 | 154,423,159 | 154,340,636 | 17 |
| | 0 | 154,756,480 | 154,933,673 | 154,845,076 | 19 |
| | 0 | 154,813,142 | 154,963,617 | 154,888,379 | 19 |
| | 0 | 155,236,093 | 155,386,538 | 155,311,315 | 21 |
| | 0 | 155,869,571 | 156,011,182 | 155,940,376 | 23 |
| | 0 | 156,245,828 | 156,422,950 | 156,334,389 | 25 |
| | 0 | 156,763,312 | 156,949,996 | 156,856,654 | 27 |
| | 0 | 156,918,444 | 157,130,858 | 157,024,651 | 28 |
| | 1 | 157,089,739 | 157,266,762 | 157,178,250 | 28 |
| A | 1 | 174,780,621 | 174,961,968 | 174,871,294 | 99 |
| | 0 | 174,960,409 | 175,130,220 | 175,045,314 | 100 |
| | 0 | 175,283,924 | 175,434,463 | 175,359,193 | 101 |
| | 0 | 175,600,401 | 175,773,331 | 175,686,866 | 102 |
| | 0 | 175,886,407 | 176,036,727 | 175,961,567 | 103 |
| | 0 | 176,108,104 | 176,295,598 | 176,201,851 | 105 |
| | 0 | 176,558,298 | 176,714,279 | 176,636,288 | 106 |
| | 0 | 177,180,236 | 177,391,475 | 177,285,855 | 109 |
| | 0 | 177,747,748 | 177,891,719 | 177,819,733 | 111 |
| L | 1 | 153,367,866 | 153,518,504 | 153,443,185 | 13 |
| | 0 | 155,275,054 | 155,425,545 | 155,350,299 | 21 |
| | 0 | 157,394,838 | 157,556,421 | 157,475,629 | 29 |
| | 0 | 159,499,529 | 159,658,201 | 159,578,865 | 38 |
| | 0 | 163,510,707 | 163,671,832 | 163,591,269 | 54 |
| | 0 | 167,491,540 | 167,680,298 | 167,585,919 | 70 |
| | 0 | 169,360,041 | 169,519,360 | 169,439,700 | 77 |
| | 0 | 171,415,729 | 171,565,827 | 171,490,778 | 85 |
| | 0 | 173,507,237 | 173,672,089 | 173,589,663 | 94 |
| | 0 | 176,530,621 | 176,711,968 | 176,621,294 | 99 |
| | 0 | 177,858,104 | 178,045,598 | 177,951,851 | 104 |
| | 0 | 179,679,177 | 179,836,567 | 179,757,872 | 119 |

Some column names are explained in the footnote of Table S2.

TABLE S4. Chromatin interactions captured by ChIA-PET between the $\alpha$-globin gene of $\alpha$-globin domain of human chr16 in two distinct cell lines (K562 and GM12878) and the rest of the domain.

| Cell line/ | $i$ | | $j$ | |
|---|---|---|---|---|
| Protein | Start(bp) | End(bp) | Start(bp) | End(bp) |
| K562/ | 228,606 | 232,911 | 101,957 | 105,248 |
| Pol II[10] | 223,848 | 234,650 | 112,365 | 122,815 |
| | 225,207 | 234,180 | 123,062 | 131,022 |
| | 228,466 | 231,601 | 140,986 | 144,385 |
| | 230,572 | 233,311 | 144,684 | 148,565 |
| | 223,675 | 234,904 | 148,901 | 176,075 |
| | 212,504 | 217,730 | 187,975 | 191,457 |
| | 228,181 | 231,216 | 193,260 | 195,050 |
| | 227,544 | 232,194 | 281,991 | 287,514 |
| | 228,369 | 231,250 | 337,052 | 340,267 |
| | 231,278 | 233,446 | 400,120 | 404,207 |
| | 227,874 | 230,293 | 415,011 | 417,686 |
| K562/ | 231,119 | 231,985 | 115,418 | 116,419 |
| CTCF[10] | 230,063 | 230,936 | 115,816 | 116,534 |
| | 231,349 | 231,926 | 118,457 | 119,027 |
| | 229,989 | 230,841 | 146,609 | 147,247 |
| | 231,126 | 232,019 | 146,696 | 147,500 |
| | 229,955 | 230,849 | 156,728 | 157,605 |
| | 231,083 | 232,073 | 156,717 | 157,821 |
| | 230,241 | 231,130 | 157,831 | 158,483 |
| | 229,997 | 230,876 | 167,564 | 168,163 |
| GM12878/ | 230,169 | 231,829 | 115,412 | 117,393 |
| CTCF[11] | 231,254 | 231,916 | 146,941 | 147,379 |
| | 233,653 | 235,304 | 157,106 | 157,280 |
| | 230,334 | 232,616 | 154,799 | 158,610 |
| | 230,238 | 232,162 | 166,032 | 168,889 |
| | 231,353 | 231,816 | 412,084 | 412,434 |

Pol II-mediated interaction, involving $\alpha$-globin genes and the rest of the domain, is absent in GM12878 cells[11]. Since CTCF-mediated interactions are mostly overlapped between K562 and GM12878 cells, the K562-specific interactions are mainly mediated by Pol II.
